# Network based multifactorial modelling of miRNA-target interactions

**DOI:** 10.1101/2020.11.16.384826

**Authors:** Selcen Ari Yuka, Alper Yilmaz

## Abstract

Competing endogenous RNA (ceRNA) regulations and crosstalk between various types of non-coding RNA in human is an important and under-explored subject. Several studies have pointed out that an alteration in miRNA:target interaction can result in unexpected changes due to indirect and complex interactions. In this paper, we defined a new network-based model that incorporates miRNA:ceRNA interactions with expression values and then calculates network-wide effects after perturbation in expression level of element(s) while utilizing miRNA interaction factors such as seed type, binding energy. We have carried out analysis of large scale miRNA:target networks from breast cancer patients. Highly perturbing genes identified by our approach coincide with breast cancer associated genes and miRNAs. Our network-based approach helps unveiling the crosstalk between node elements in miRNA:target network where abundance of targets leading to sponge effect is taken into account. The model has potential to reveal unforeseen and unpredicted regulations which are only evident when considered in network context. Our tool is scalable and can be plugged in with emerging miRNA effectors such as circRNAs, lncRNAs and available as R package ceRNAnet-sim https://www.bioconductor.org/packages/release/bioc/html/ceRNAnetsim.html.

## Introduction

MicroRNAs (miRNAs) are a family of short non-coding RNAs which are key regulator of gene expression through various post-transcriptional mechanisms (1). Although the mechanisms by which miRNA effects are not fully understood, miRNAs predominantly repress their targets. Repressive activities of miRNAs vary depending on many factors that are significant to miRNA:target interactions. These factors include miRNA:target binding energy, binding location in target sequence, base pairing types between miRNA and target, abundance of miRNAs and targets (2). For example, a proteomics study have investigated the importance of seed pairing type between miRNAs and their targets and target site location and proposed that the characteristics of binding between miRNA and target drastically affect miRNA efficiency (3). Similarly, another study revealed that affinity is correlated with seed pairing of miRNA:target pairs and suggested that affinity is correlated with length of canonical seed base pairing (4). Binding energies of miRNA:target complexes vary based on nucleotide context and determine folding stability of miRNA:target complex (5). It has been demonstrated that the binding energy between miRNA and target indicates stability or affinity of complex (6, 7) and does not directly determine repressive activity of miRNA (5). Early studies have argued that 2-8 nt sequence located in 5’end of miRNA, known as seed, bind to specific sequence located in 3’UTR of its target (8, 9). In recent studies, it has been shown that miRNAs can interact with targets via sequences located in various regions such as 5’UTR or CDS (6, 10, 11). These studies also showed that binding location either dictates affinity of miRNA:target interaction or affects level of target degradation. It has been shown that miRNAs exhibit repressive activity via 6-8 nt long sequence that is perfectly complementary with seed region of their targets (2, 12). On the other hand, some researchers have reported that seed sequence of miRNA can have mismatches or bulged/wobble nucleotides (13). On top of all these factors, abundance of miRNAs and targets and miRNA:target ratio in cells predominantly affect efficiency of miRNA:target interaction (4, 14, 15).

As it is possible for miRNAs to suppress multiple targets, an individual mRNA molecule can also be targeted by multiple miRNAs. In that case, the targeted mRNAs exhibit competitor behavior, hypothesized as competing endogenous RNAs (ceRNAs) against their miRNAs (16, 17). Briefly, (16) have explained the ceRNA hypothesis as disturbance of the other target when one of the targets was perturbed with expression change on a steady-state system that included one miRNA and two targets (16). Regarding interaction between miRNAs and their targets in a cell, explaining and predicting aftermath of an individual perturbation is difficult due to complexity of interactions. Various computational and experimental studies have tackled the problem of unraveling ceRNA:miRNA interactions. For instance, when abundance of one of the targets of miR-122 increases, the expression levels of remaining targets also slightly increase as a result of decreasing repressive activity of miR-122 on remaining targets (15). (4) have developed a mathematical model for changes on total target pool concentration after grouping targets according to affinity and demonstrated that miRNA activity correlated with affinity between miRNA and target (4). Cooperative efficiency of miRNAs as well as competitor behaviors of targets were also studied and it has been demonstrated to be crucial for regulating available mRNA levels of targets (18). MiRNA:target interactions have been modeled as stoichiometric and catalytic mechanisms and (19) have recommended handling models in network context. Models that can explain miRNA target interactions through topological features has been applied to bipartite networks considering direct interactions only (20) or considering both direct and indiract interactions (21). Through common miRNAs and genes, all miRNAs and targets in the network were shown to interact with each other in bipartite fashion. More recently, an approach to detect ceRNA pairs by using the miRNA expression, gene expression and common miRNAs between gene targets was developed (22) which is effective in analyzing genes through miRNAs. The authors have concluded that existing miRNA based approaches may not be suitable for understanding regulations of ceRNA interactions. Additionally, miRmapper package (23) utilizes an adjacency matrix to associate miRNAs using differentially expressed genes and identifies significant nodes using topological properties of network.

## Results

### Network with miRNA:ceRNA expression level

We have developed a network-based approach to assess effects of expression level changes in ceRNA regulation. The basic mode of miRNA repression activity has been established based on miRNA and target abundance in various researches (14, 15). Our approach can easily calculate effects of expression changes to whole network when expression level is only available factor. In Sample network given in Figure 1 and Table 1, after an increase in expression level of a gene (G2), expression values of other genes also changed due to redistribution of miRNA among its targets. Previous studies have shown that if abundance of a gene increases in a ceRNA network, expression levels of remaining targets are affected due to shared targets of multiple miRNAs (24– 26). Accordingly, effect of perturbation in a single gene can spread though the whole miRNA:target network.

**Fig. 1.**
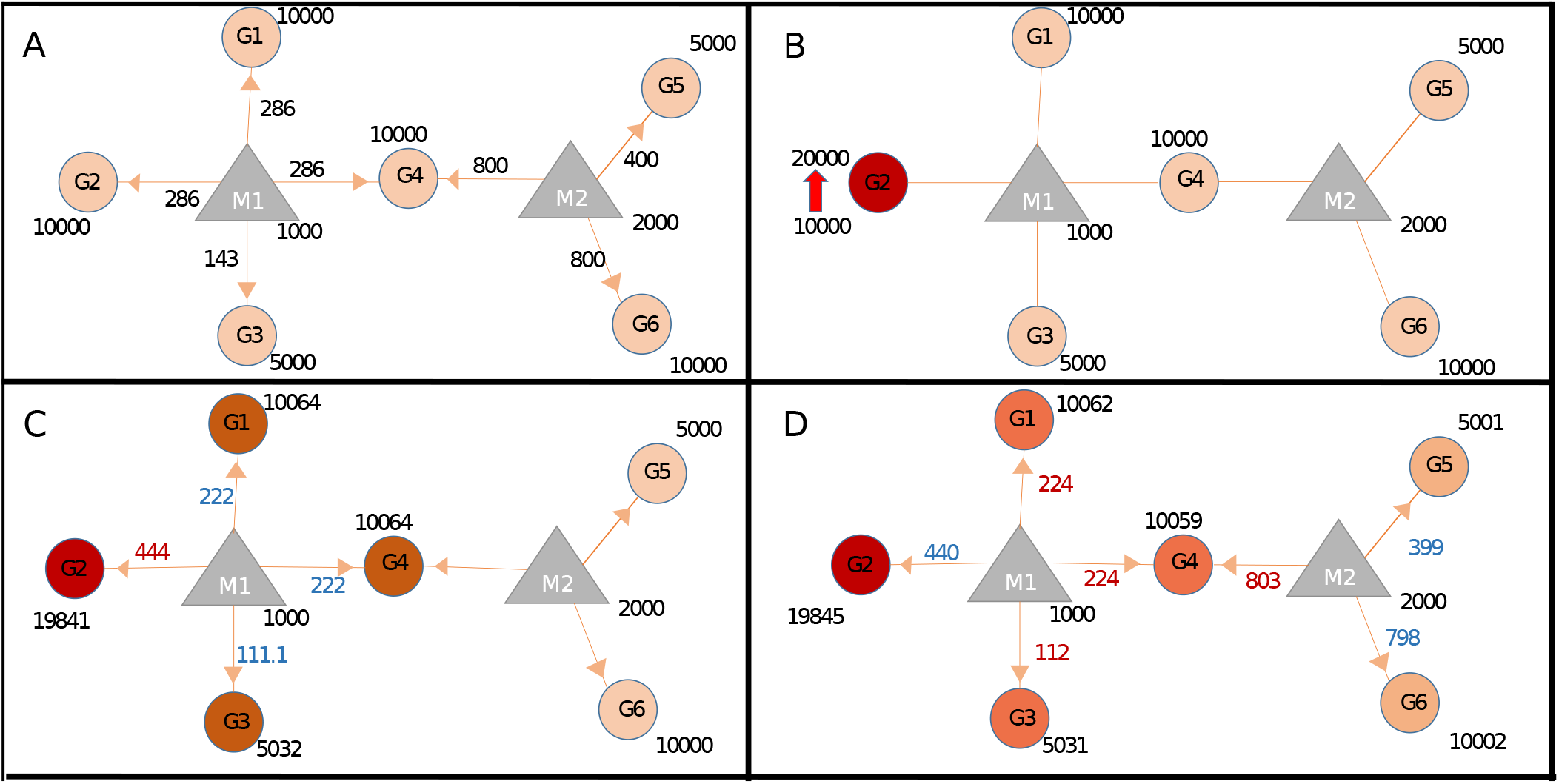
Schematic presentation of mechanism of network based model. (**A**) In steady state, miRNAs (M: triangles) repress targets (G: circles) according to proportion of target expression level. (**B**) Two fold increase in transcript level of Gene2 (G2) acts as trigger (shown with thick red arrow). (**C**) Distribution of miRNA1 (M1) changes due to increased distribution of M1 over G2 (286 to 444 units) hence decreasing its distribution to lower levels for G1, G4 (both from 286 to 222) and G3 (from 143 to 111.1), updated distribution levels are shown in blue and red numbers. Due to less miRNA targeting, genes G1, G3 and G4 show increased levels of availability, from 10000 to 10064 or from 5000 to 5032. (**D**) The change at expression of common target, G4 which pulls more M2 hence decreasing availabilty of M2 for both G5 and G6, consequently levels of G5 and G6 increase due to decreased repression by M2. Expression values are rounded to integers for simplicity. Values on edges indicate initial distribution, red and values indicate increase and decrease in distribution, respectively. Shades of circles indicate different levels of increase.

**Table 1.**
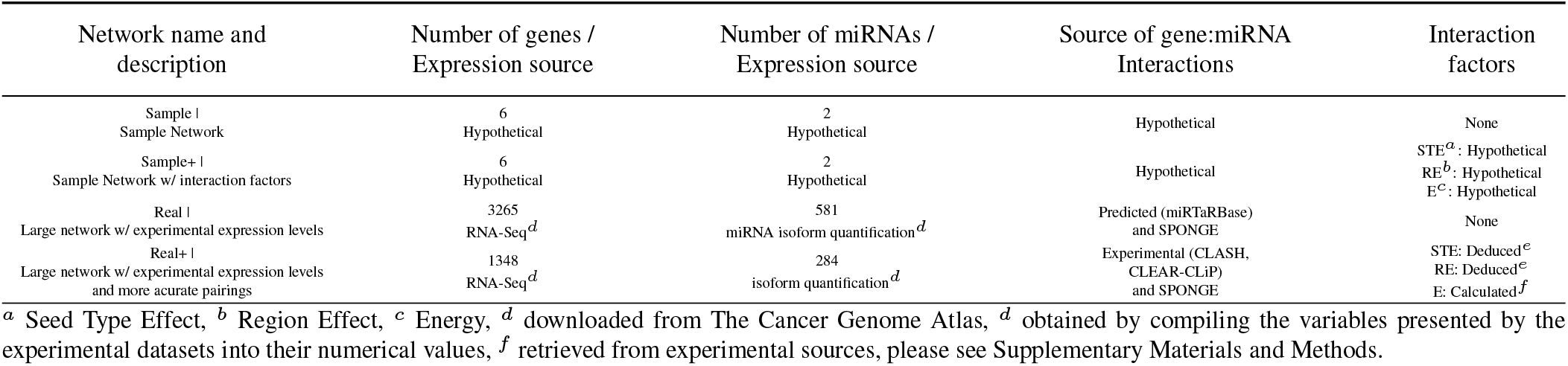
Summary of tables used in this study.

Genes targeted by multiple miRNAs act as a trigger for adjacent local neighborhood of targeting miRNAs, causing changes in expression levels of genes outside the local neighborhood of initial trigger gene. Therefore, primary expression change in gene (G2) causes changes in other group of genes (G5 and G6) even though initial trigger gene (G2) and genes in other group are not targeted by common miRNA (Figure 1D). In addition, as shown by an earlier ceRNA hypothesis model (16), after the increase of gene expression level of G2, the miRNA that is found in the same group (M1) becomes less repressive on its remaining targets (G1, G3 and G4). It’s important to note that the changes in gene expression levels will have more pronounced effect if miRNA:target ratio is high, i.e., more miRNA available per target, which was reported in previous findings (4, 14, 15). Our findings showed that more repressive activity occurred on genes targeted by more than one miRNAs.

### ceRNA:Target networks with interaction factors

Our approach can calculate miRNA reppression activity more accurately, by integrating seed type, region and energy parameters. To demonstrate this capability, we generated Sample+ network, summarized in Table 1 and Table S1, for further calculations. Proportional distribution of miRNA on targets is determined in accordance with equations Eq. (1) and Eq. (2), when the model run based on expression values of miRNAs and ceRNAs (genes). In that case, the system reaches a steady-state shown in Figure 2A. However, if interaction factors are taken into account, specified interaction factors (i.e, binding factors such as seed type and energy shown in Table S1) affect distribution of miRNA on targets as calculated in equation Eq. (3) and described in Figure S5C. After affinity based proportional distribution of miRNA expression, degradation factor is considered to specify amount of repressive miRNA in pairs because not all miRNA:target binding events result in degradation (Figure S5D). For instance, proportional distribution of G1:M1 interaction in Figure 1A is calculated higher than the same pair (G1:M1) in Sample+ network (Figure S5C) (286 vs. 6.58). On the other hand, calculations with interaction paramaters could cause increasing miRNA repression activity like in miRNA2:Gene4 interaction (Figure 2A). When expression of Gene2 (G2) increased (Figure 2B and Figure S7), expression values of all genes also changed at various levels because of contribution of efficiency factors (Figure 2B-D and Figure S7-8). These results suggest that the unexplained regulations in miRNA:target interactions can be revealed by converting the interaction parameters into numerical expressions.

**Fig. 2.**
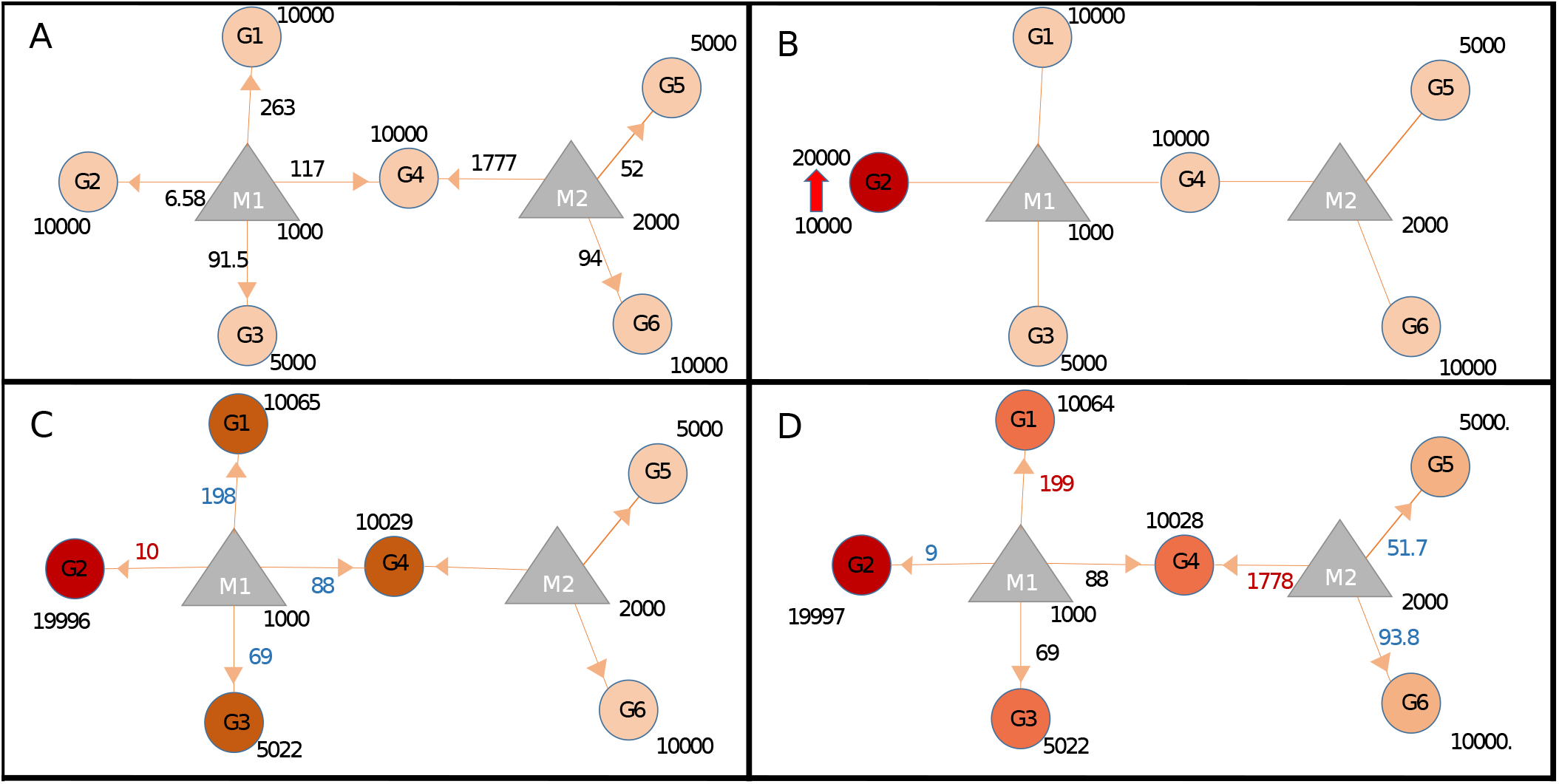
Target regulations with interaction parameters. (**A**) In the steady-state the repression activity of miRNAs on the targets after binding and repression efficiency. (**B**) The changes the repression activities after increasing of G2 expression. (**C**) Perturbation of primary neighborhoods of M1 miRNA (M1 miRNA group). (**D**) Regulation of gene expression of other gene group via triggering target (common target between M1 and M2)

When the factors were taken into account in the system, miRNA efficiencies varied as shown in Figure 2A (also refer to Figure S6). Although the miRNA:target expression ratios in steady-state were same in comparison with the Sample dataset, efficiency of binding and repression have changed. On the other hand, changes in expression levels of G4 and G1 differ although they have same expression levels at initial conditions. This is due to fact that G4 is targeted by two miRNAs, M1 and M2 (Figure 2D) and additional interaction parameters.

When Sample+ network is triggered with two fold increase in expression level of common target (G4), more prominent changes were observed in gene expression levels in network compared to changes caused by G2 over-expression (Figure S9). Furthermore, change in expression level of target gene that has strong miRNA repression efficiency resulted in evident perturbation in interaction network. On the other hand, it was observed that Gene2 was weakly affected by changes in expression level of Gene4 due weak interaction factors. Finally, our function screens each gene in the network for their perturbation efficiencies, while considering various ceRNA interaction parameters. When we applied the method on the Sample+ dataset, Gene4 has been found to be the most effective node in terms of number of perturbed elements and miRNA2 (M2) has been found to be causing the highest mean in expression changes (see Section 1 in Supplementary Materials and Methods).

### Compiling Realistic Network and Identifying Nodes That Cause Widespread Perturbation

For large scale analysis, we constructed the Real network with miRNA:target pairs from miRTarBase comprising of 3265 genes and 581 miRNAs (Table 1). Many instances of the Real network were constructed by overlaying expression data for each patient. We constructed 174 networks from 87 patients who had both normal and cancer expression data available. Subsequently, perturbation efficiency of each node in each network was calculated. As a result, 70 of 3265 genes and 27 of 581 miRNAs were found to have high number of perturbing node in both tissues, normal and cancer (Figure 3A). It has been observed that 29 genes and 1 miRNA had tumor tissue specific perturbing activity. Additionally, in normal tissue samples of these 87 patients, 46 genes and 4 miRNA have showed robust perturbation efficiency. Please note that tumor-specific perturbing genes not necessarily exhibit differential expression between normal and cancer tissues (shown at Figure S11).

**Fig. 3.**
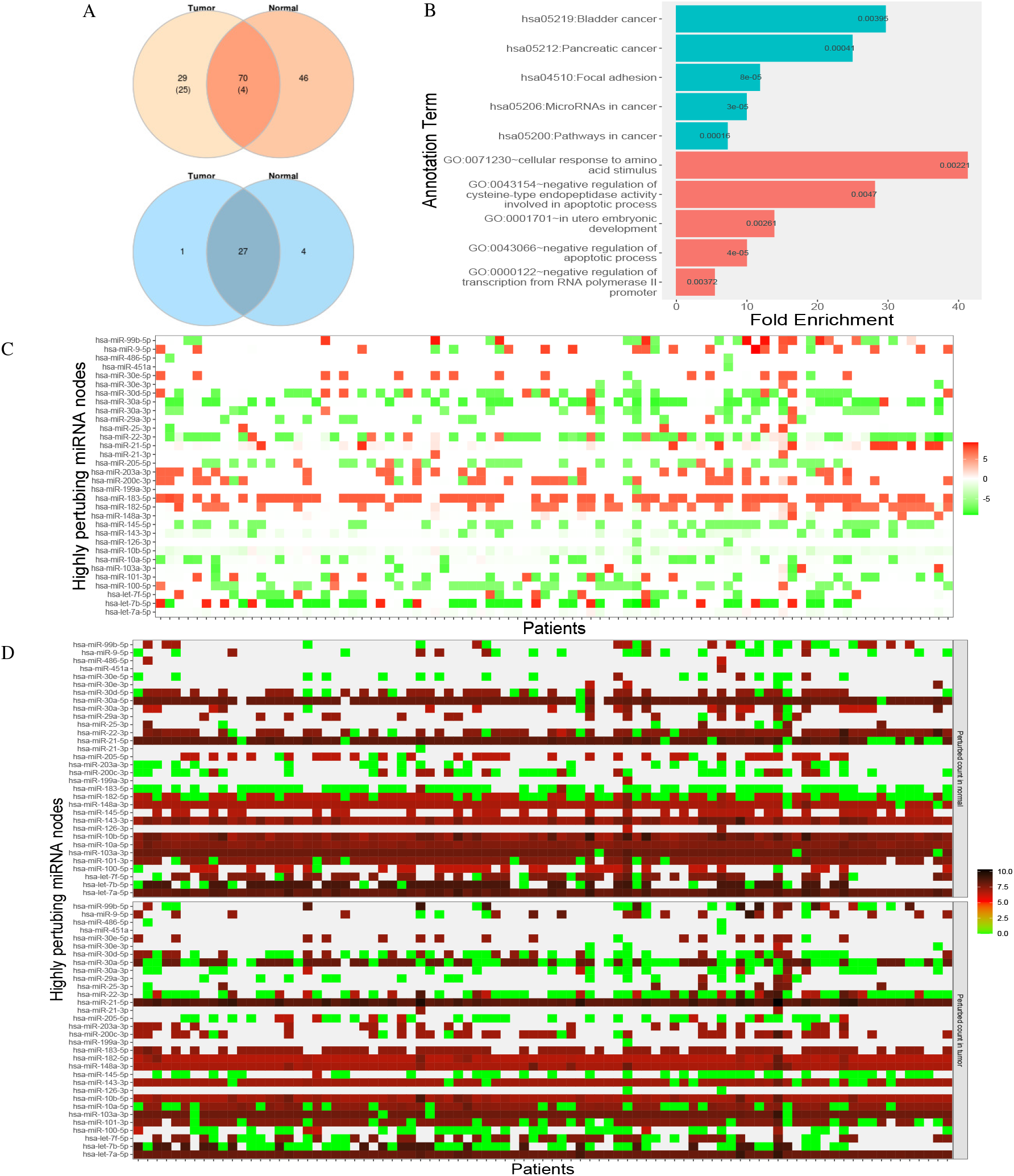
Analysis of highly perturbing genes and perturbed node counts for all miRNAs in Real network. **(A)** Number of highly perturbing nodes (genes in orange, miRNAs in blue) with miRNA:gene target pairs, 99 perturbing genes were analyzed at DisGeNet. Number of breast cancer associated genes are indicated in parenthesis. DisGeNet analysis was not performed for normal specific perturbing genes. **(B)** Top five enriched KEGG pathway (teal) and Gene Ontology (red) terms using all tumor-specific highly perturbing genes, **(C)** Comparison of perturbed node counts in tumor and normal tissues for each miRNA. Represented as log2 of affected node count ratio between tumor and normal samples, **(D)** Affected node number of critical miRNAs in tumor and normal tissues. Represented as log2(affected node count).

Our findings accord with results from different tools and databases. Considerable number of these genes, exactly 44, were enriched (p-value < 0.05) in critical pathways in cancer such as PI3K-Akt signaling pathway (4.69 fold enrichment, p-value 2.04e-08), proteoglycans in cancer (6.07 fold, p-value 1.03e-07), FoxO signaling pathway (6.64 fold, p-value 4.55e-06). It has been observed that 122 of these genes have enriched in biological processes and molecular functions, including negative regulation of apoptotic process (GO:0043066, 4.75 fold and p-value 5.15e-07), cellular response to epidermal growth factor stimulus (GO:0071364, 23.12 fold and p-value 5.33e-06), cadherin binding involved in cell-cell adhesion (GO:0098641, 8.25 fold, p-value 1.35e-11), transcription factor binding (GO:0008134, 5.32 fold, p-value 1.54e-05). Interestingly, tumor specific perturbing genes show enrichment in cancer associated pathways and biological processes (Figure 3B) while normal tissue specific perturbing genes were not enriched significantly in same pathways or processes. Enriched perturbing genes have been illustrated in perturbing node network in Figure 4. While four of top five enriched KEGG pathways were evidently associated with cancer, remaining pathway (focal adhesion) has been shown to be associated with adhesion, migration and invasion in breast cancer (27). Additionally, regulation of amino acid metabolism (28) and apoptotic processes which play significant role in tumor progression, were found enriched by tumor-specific perturbing genes in these process (Figure 3B). Also, negative regulation of transcription regulation from RNA polymerase II is important in manipulating tumor microenvironment and communications of cancer cells. In utero embryonic development process (GO:0001701) was found to be enriched in our results and it was shown to play role in breast cancer metastasis and mammary development in utero (29).

**Fig. 4.**
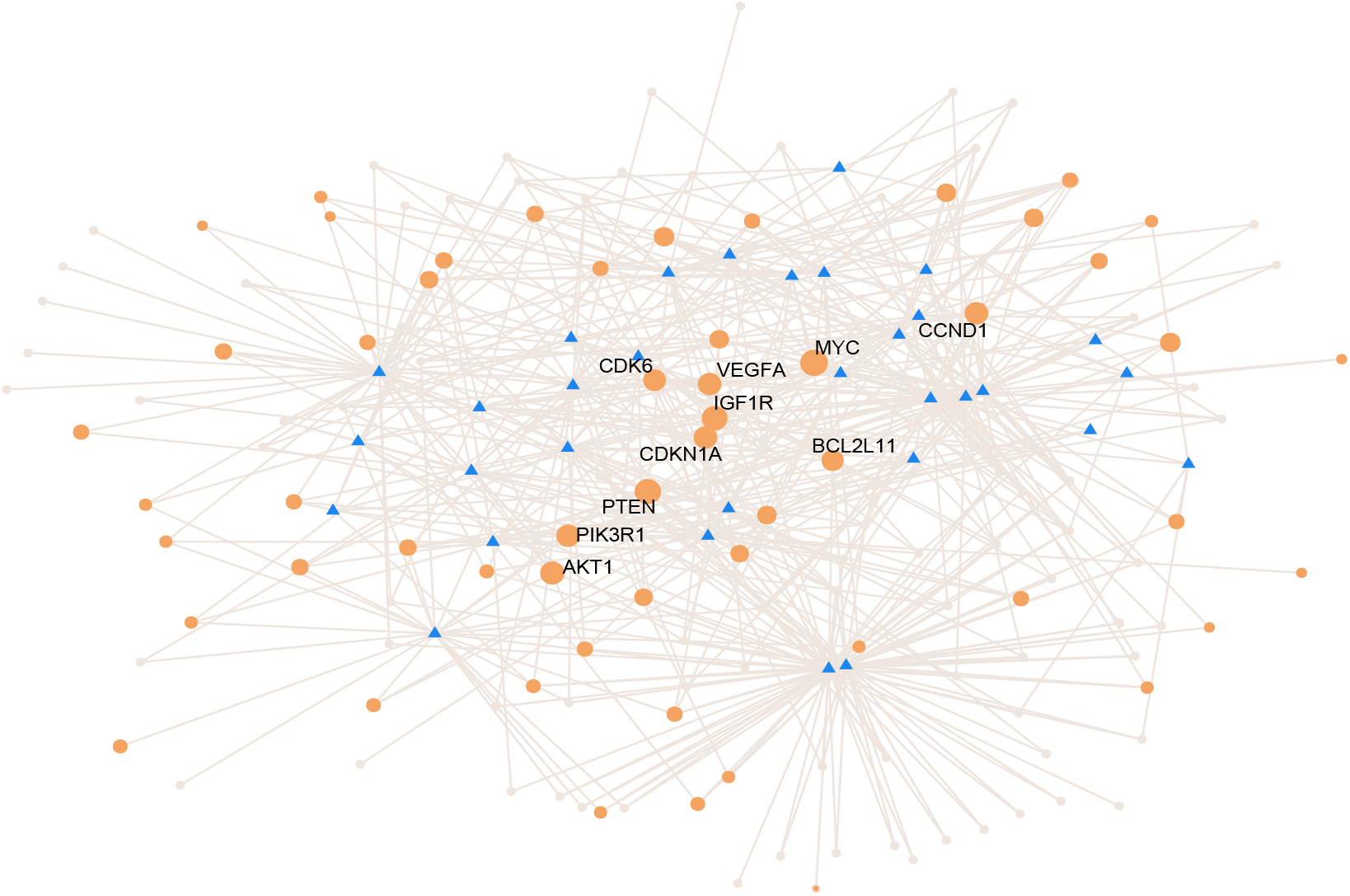
Network of miRNA:genes in enriched KEGG pathways and GO terms. Highly perturbing genes (circles) and miRNAs (triangles) in Real network have been illustrated. Tumor-specific genes enriched in top 5 KEGG or GO terms are colored in orange. Top ten genes that have highest degree centrality are labeled.

DisGeNet platform (30), a database listing critical genes for numerous diseases, was used to analyze critical genes in our findings. We found that 29 of 99 genes in tumor tissue have breast cancer disease association score greater than 0.1, with 6.95e-08 hypergeometric p-value and 2.9 fold enrichment.

In order to characterize highly perturbing miRNAs they were examined via HMDD, a database compiling miRNA and disease relationships regardless of isoform differences (31). While 481 of 1208 miRNAs are associated with breast cancer in whole HMDD database, nearly all highly perturbing miRNAs (28 of 29) in our study were denoted as breastb cancer associated (2.42 fold enrichment with p-value 7.12e-11). 28 high perturbing miRNAs have been associated with breast cancer, 20 of them were attributed with causality. The remaining non-breast cancer associated miRNA, miR-99b, may play a regulatory role on proliferation and migration processes in breast cancer by affecting the TGF-*β* signaling pathway (32).

We also evaluated miRNA significance by comparing perturbing node counts for each miRNA across all samples (Figure 3 C and D). For example, miR-30a-5p isoform which suppresses proliferation, migration and tumor growth (33) has been observed as highly effective in almost all normal tissue samples but not in all tumor tissue samples. As another example, miR-183 which is commonly up-regulated in tumor samples and one of the key regulators of metastatic process in breast cancer tissues (34), exhibited diverse perturbing efficiencies between normal and tumor tissues in our analysis. miR-182-5p and miR-21-5p have been found as highly efficient in tumor tissues, but not in all normal tissues. These results indicate that our approach suggests a new perspective to ceRNA network analysis and can contribute to disease-therapy studies with ceRNA network topology dependent method.

### Detecting Perturbation Efficiency of Nodes in Presence of Interaction Parameters

For more accurate analysis we generated a network with experimental data. The interactions in the Real+ network originated from CLEAR-CLiP and CLASH datasets as opposed to the Real network which is comprised of predicted miRNA:target interactions. 1348 genes and 284 miRNAs associated with these genes were compiled as the Real+ network, see Table 1. While significant factors such as energy, seed type, and location of seed on the target sequence in miRNA target interaction in the template network are same for each patient dataset, expression levels of miRNA and gene differ in each dataset. We found the perturbation efficiencies for each node in each patient by simulating perturbation in expression level of every miRNA and gene in the network.

By fitting mixed distribution model to simulated networks with mean expression values from random patient data sets, a node is defined as “perturbing node” if it effects at least 216 nodes in at least 18 samples. It has been observed that 475 of 1348 genes were with high perturbation efficiency in normal and tumor tissue samples. While 12 of these genes were highly effective in tumor tissues specifically, no genes were detected showing normal tissue-specific effect.

On the other hand, 83 of 284 miRNAs were found to be high perturbing node in both tissues and only one miRNA per tissue group have been found specific to that tissue group. KEGG or GO term enrichment analysis of 475 perturbing genes by DAVID tool (35, 36) (Figure S10) showed that 60 of these genes were enriched in signal pathways like FoxO signaling pathway (p-value 0.0058 and 2.78 fold enrichment). Also 399 of perturbing genes were significantly enriched in cancer associated biological process and molecular functions such as cadherin binding involved in cell-cell adhesion (GO:0098641 with p-value 1.19e-08 and 3.68 fold enrichment), cell-cell adhesion (GO:0098609 with p-value 5.30e-08 and 3.65 fold enrichment), microtubule cytoskeleton organization (GO:0000226 with 5.23e-04 p-value and 4.82 fold enrichment), negative regulation of Ras protein signal transduction (GO:0046580 with p-value 3.80e-03 and 7.61 fold enrichment). When we analyze the critical genes associated with breast cancer from the DisGeNET database (30), 32 of the disease related genes with a score greater than 0.1 were observed (1.50 fold enrichment with p-value 0.0051).

Regardless of the isoform difference, 67 of 79 miRNAs (2.13 fold enrichment with p-value 2.05e-17) have been associated with breast cancer in the HMDD database (Version 3.2) (31) and 40 of these miRNAs reported as having potential to cause breast cancer. Remaining 12 miRNAs not associated with breast cancer in HMDD were explored further in literature. There are numerous studies that associate breast cancer with miR-28 (37), miR-1287 (38), miR-3065 (39), miR-500a and miR-99b (32). They have the potential to be used for biomarker, therapeutic or regulatory purposes. Additionally, miR-501, miR-532 and miR-589-3p, which have perturbation efficiency in tumor tissue, were not associated with breast cancer in HMDD. Several researches about miR-589-5p association with drug response (41) and abnormal regulation of miR-532-5p (42) in breast cancer were reported. miR-577 have been reported as potential target for breast cancer therapy in an earlier study due to its function as suppressor of metastasis which induced by epithelial-mesenchymal transition (43). On the other hand, at the time of writing this study there were no reports that directly associate hsa-mir-2116, miR-3127 and miR-501-3p with breast cancer.

Compared to simulations made with networks constructed with miRTarBase (44) data set, results from Real+ network showed that high number of genes/miRNAs perturbed large number of nodes. This finding emphasizes the importance of integrating miRNA:target interaction parameters as they profoundly effect the proportional distribution of miRNA expression. As more miRNA:gene interaction parameters become available in future studies by novel experimental methods, our method has the potential to integrate them and thus providing more accurate depiction of ceRNA network and its consequences.

### Evaluation of Performance and Runtime

Our tool can successfully simulate perturbations in large network despite complex behaviors where reaching the steady-state is a challenge. Simulations show that change in expression level of single gene has potential to affect whole network, perturbing many distant nodes. These observations are in accordance with competing endogenous RNA hypothesis where genes targeted with many common miRNAs subsequently transmit perturbation to neighboring groups.

## Discussion

Compared to earlier attempts of analyzing miRNA:target interactions, our approach can operate on large-scale networks while integrating interaction parameters on top of expression level data. In order to process large scale networks, steady state approach was taken in our approach. Although kinetic modeling is more appropriate and accurate for modeling expression levels, its calculation for large networks is impractical. Thus we used steady-state approach to process large networks. Transcription, degradation or binding rates of miRNAs or mRNAs are ignored during calculations. Although miRNAs are known to be stable, the transcription and degradation rates of miRNAs change depending on cellular conditions (45). Additionally, other regulation parameters such as gene-gene interactions and activations by transcription factors are ignored as well.

Although we considered only mRNAs as ceRNAs in our approach, other non-coding RNA (ncRNA) types are known to compete for miRNA interaction, consequently acting as sponges for miRNAs. Such ncRNAs play role in regulation of crucial genes. For instance, critical genes extracted from lncRNA-miRNA-mRNA interactions overlap with genes important for cancer (46). More specifically, involvement of lncRNA-miRNA-mRNA interactions with breast cancer (47, 48) and involvement of circRNA-miRNA-mRNA interactions with cervical cancer (49) has been shown in recent studies. Our calculations can easily integrate expression level and miRNA targeting data for various ncRNAs as they become available. Our approach is not only capable of integrating various players, but also various parameters. Yet to be discovered interaction parameters affecting binding and post-binding repression by miRNAs can easily be integrated into calculations. For instance, recent findings show that not all miRNA:target binding events result in functional repression (50). As soon as its data becomes available such a parameter can be included in our calculations.

It’s noteworthy that our findings about tumor specific perturbing genes are derived from ceRNA expression levels in context of network topology not from differential expression analysis. Some of the genes that are detected as perturbing genes have comparable expression levels between normal and tumor samples (Figure S11).

Differential expression analysis is commonly used for identifying genes or miRNAs that are important between two conditions, usually normal vs disease. Our approach has potential to reveal critical genes even though their expression does not change. By harnessing the complexity of ceRNA network, we observe that a gene or miRNA might become critical if the expression level of gene(s) in its local neighborhood changes. As an example, we evaluated miRNA significance by comparing perturbed node counts for each miRNA across all samples (Figure 3 C and D). Although miR-30a-5p has comparable expression between normal and tumor samples (Figure S12), it was shown to be effective in normal sample only. Considering the fact that miR-30a-5p suppresses proliferation, migration and tumor growth (33), our finding might explain how miR-30a-5p loses it suppression without changing its expression level.

In small datasets, finding perturbation efficiency of each element, determination of iteration number to reach steady-state have relatively short runtimes (51). However, networks integrating all miRNA:target interactions require immense calculation which increases the runtime significantly. Although we took advantage from parallel processing capabilities of various R packages, the runtime for perturbation efficiency calculation is still longer than anticipated.

## Conclusion

Our study has ability to integrate expression values of mRNA and miRNAs along with their interaction parameters into a life-size miRNA:target network. Consequences of a perturbation can be simulated for a node in whole network. Moreover, perturbation analysis can be performed for each node, which reveals nodes with high perturbation efficiency. Considering the competition between mRNAs provides more accurate analysis of ceRNA networks. The ceRNAnetsim package is extensible by integrating various interaction parameters as more experimental data becomes available. Moreover, additional players in ceRNA network such as non-coding RNAs can be integrated into calculations as soon as their expression levels and miRNA targeting data becomes available. Our package is able to reveal critical genes which are not discovered by conventional approaches such as differential gene expression analysis. Consequently, our package may help researchers tackle complex interactions in ceRNA networks with a novel approach, leading to better understanding and predictions of abnormal regulations and pathways underlying diseases or conditions.

## Methods and Materials

### Construction of miRNA:target network

For Sample network minimum required information, expression level of genes and miRNAs, was used. The Sample+ network had additional columns on top of Sample network (Table 1). miRNA:gene pairs from miRTaRBase, expression values of miRNA and genes from TCGA were fed into SPONGE for sparse partial correlation analysis (22). Resulting miRNA:gene pairs were used to construct the Real network. In comparison, edge data of the Real+ network was retrieved from CLASH and CLEAR-CLiP (6, 11). Additional parameters for the Real+ network were curated from literature (see section 2 in Supplementary Materials and Methods). Table 1 summarizes each network. While constructing custom networks, users can plug additional custom parameter(s) and indicate its effect (degradation, affinity or both) to be considered.

### Triggering Perturbation and Subsequent Calculations

Initially, the network, Sample network at Table 1, is assumed in steady-state (Figure 1A and Figure S2) condition and needs least one trigger for initiating calculations. The trigger can be a change in expression level of one or more genes (Figure 1B and Figure S3). After trigger, the network undergoes iterative cycle of calculations at each of which distribution of miRNA in local neighborhood is recalculated (Figure 1C). Based on new miRNA distribution, expression level of each node (i.e. ceRNA) is updated (Figure S4). Due to common targeted elements, the change in one neighborhood spreads to other neighborhoods (Figure 1D), consequently have potential to affect whole network due to “ripple effect”.

During calculations, following assumptions were adopted; 1) Transcription and degradation rates of miRNAs are steady and equal. 2) All available miRNAs are recycled as in miRNA:ceRNA binding, target is degraded and miRNA is unaffected. 3) ceRNA targets also have stable transcription and degradation rates and these rates are equal. The repression efficiency of a miRNA on the individual target (*Eff*_*ij*_) is calculated according to equation Eq. (1); where miRNA expression (*C*_*j*_) in local neighborhood is distributed among targets using individual gene expression levels (*C*_*i*_), *k* number genes targeted by *jth* miRNA. For the genes targeted by multiple miRNAs, *n* number of targeting miRNAs for *ith* gene, cooperative activity of miRNAs on a target gene, *R*_*i*_, is calculated by summing repression activity of each miRNA Equation Eq. (2).

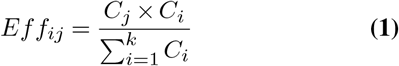

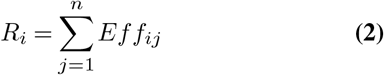

### Multifactorial calculations in miRNA:target network

Interactions between miRNAs and their targets can be affected from various factors. So, our model integrates multiple factors when calculating overall miRNA activity. We classified factors into two categories. Binding factors determine interaction between miRNA and target and they alter amount of miRNA sequestered to target. Efficiency factors dictate degradation efficiency of sequestered miRNA on its target. In other words, binding factors exert their influence before or during binding, efficiency factors exert their influence after binding.

From the literature, binding free energy (5, 6), seed type (52) and binding region (e.g, 5’UTR, 3’UTR, CDS) (6, 10) data has been retrieved and plugged into Real+ network. For Sample+ at Table 1 network we used arbitrary values.

The normalized values of factors are used to determine binding activity and miRNA efficiency on targets (Figure S5) using equations 3 and 4. Binding affinities (activity, *Eff*) of miRNAs on each individual gene are calculated as shown in equation Eq. (3) where *STE*, seed type effect; *RE*, Region Effect; *E*, Energy; ′, normalized values of these factors, *k* number genes targeted by *jth* miRNA; *C*_*j*_, miRNA expression; *C*_*i*_, individual gene expression (Figure S5C).

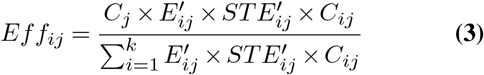

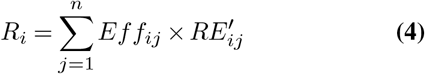

Not all miRNA:target binding events result in degradation of target. The degradation of target by bound miRNA depends on efficiency factors such as binding region. Exact repression efficiency of miRNA is calculated according to equation Eq. (4) (Figure S5D);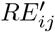, normalized values of region efficiency coefficient between miRNA and gene, *n* number of targeting miRNAs for *ith* gene, *R*_*i*_ cooperative activity of miRNAs on a target gene. The cooperative repression activity of miRNAs to their common targets is figured out as shown in Figure S5E.

### Analysis of Real Breast Cancer Networks and Determining Node Efficiencies

Expression levels of miRNA and genes in tumor and normal tissues of 87 patients are retrieved from TCGA; https://www.cancer.gov/tcga. We used sparse partial correlation method, SPONGE, offered by the (22) to filter out genes with weak association. SPONGE R package (53) was used on log transformed miRNA and gene expression datasets of breast cancer patients and selected genes with p-adj less than 0.2, as suggested by authors. For Real network, predicted miRNA:target interactions from miRTarBase (44) and for Real+ network, experimental miRNA:target dataset (i.e. CLEAR-CLIP and CLASH datasets) (6, 11) have been used as binary matrix for correlation analysis.

Our package has functions to calculate perturbed count and perturbation efficiency for each gene by using each gene as a trigger. Perturbed count is number of affected nodes perturbation efficiency is defined as mean of percent change in expression level of perturbed nodes. Through parallel processing, we evaluated the perturbation efficiencies in Real and Real+ networks of all nodes in breast cancer tissue samples. Throughout the study we used two fold increase in expression level as trigger. Please note that two fold decrease as a trigger gave same results. Perturbing nodes that show high efficiency in tumor and normal tissues were analyzed further (KEGG pathway, Gene Ontology etc.).

### Calculating Threshold Values for Affected Nodes and Patients

We used mixed model to fit distribution of perturbation efficiencies of nodes in patient data sets and to determine appropriate cut-off values for number of perturbed nodes and number of patients (54). To test accuracy of cut-off value, we performed simulations on randomly selected nodes with mean miRNA and gene expression values of random 10 patient’s data sets. We considered that nodes which have perturbation efficiency at least 1 patient with 78 perturbed nodes are effective in Real network of samples. Based on results from mixed distribution model, nodes that perturbed at least 216 nodes in 18 samples were considered as perturbing node in Real+ network.

## ACKNOWLEDGEMENTS

The numerical calculations reported in this paper were partially performed at TUBITAK ULAKBIM, High Performance and Grid Computing Center (TRUBA resources).

## Bibliography

1. Julius Brennecke, Alexander Stark, Robert B Russell, and Stephen M Cohen. Principles of microrna–target recognition. PLoS biology, 3(3):e85, 2005.

2. Andrew Grimson, Kyle Kai-How Farh, Wendy K. Johnston, Philip Garrett-Engele, Lee P. Lim, and David P. Bartel. MicroRNA Targeting Specificity in Mammals: Determinants beyond Seed Pairing. Molecular Cell, 27(1):91–105, July 2007. ISSN10972765. doi: 10.1016/j.molcel.2007.06.017.

3. Wenlong Xu, Zixing Wang, and Yin Liu. The Characterization of microRNA-Mediated Gene Regulation as Impacted by Both Target Site Location and Seed Match Type. PLoS ONE, 9 (9):e108260, September 2014. ISSN 1932-6203. doi: 10.1371/journal.pone.0108260.

4. Andrew D. Bosson, Jesse R. Zamudio, and Phillip A. Sharp. Endogenous miRNA and Target Concentrations Determine Susceptibility to Potential ceRNA Competition. Molecular Cell, 56(3):347–359, November 2014. ISSN 10972765. doi: 10.1016/j.molcel.2014.09.018.

5. Song Cao and Shi-Jie Chen. Predicting kissing interactions in microRNA–target complex and assessment of microRNA activity. Nucleic Acids Research, 40(10):4681–4690, May 2012. ISSN 1362-4962, 0305-1048. doi: 10.1093/nar/gks052.

6. Aleksandra Helwak, Grzegorz Kudla, Tatiana Dudnakova, and David Tollervey. Mapping the Human miRNA Interactome by CLASH Reveals Frequent Noncanonical Binding. Cell, 153 (3):654–665, April 2013. ISSN 00928674. doi: 10.1016/j.cell.2013.03.043.

7. Jeremie Breda, Andrzej J. Rzepiela, Rafal Gumienny, Erik van Nimwegen, and Mihaela Zavolan. Quantifying the strength of miRNA–target interactions. Methods, 85:90–99, September 2015. ISSN 10462023. doi: 10.1016/j.ymeth.2015.04.012.

8. David P Bartel. MicroRNAs. Cell, 116(2):281–297, January 2004. ISSN 00928674. doi: 10.1016/S0092-8674(04)00045-5.

9. Benjamin P. Lewis, Christopher B. Burge, and David P. Bartel. Conserved Seed Pairing, Often Flanked by Adenosines, Indicates that Thousands of Human Genes are MicroRNA Targets. Cell, 120(1):15–20, January 2005. ISSN 00928674. doi: 10.1016/j.cell.2004.12.035.

10. J. Hausser, A. P. Syed, B. Bilen, and M. Zavolan. Analysis of CDS-located miRNA target sites suggests that they can effectively inhibit translation. Genome Research, 23(4):604–615, April 2013. ISSN 1088-9051. doi: 10.1101/gr.139758.112.

11. Michael J. Moore, Troels K. H. Scheel, Joseph M. Luna, Christopher Y. Park, John J. Fak, Eiko Nishiuchi, Charles M. Rice, and Robert B. Darnell. miRNA-target chimeras reveal miRNA 3’-end pairing as a major determinant of Argonaute target specificity. Nature Communications, 6:8864, November 2015. ISSN 2041-1723. doi: 10.1038/ncomms9864.

12. David P. Bartel. MicroRNAs: Target Recognition and Regulatory Functions. Cell, 136(2): 215–233, January 2009. ISSN 00928674. doi: 10.1016/j.cell.2009.01.002.

13. Sung Wook Chi, Gregory J Hannon, and Robert B Darnell. An alternative mode of microrna target recognition. Nature structural & molecular biology, 19(3):321, 2012.

14. Aaron Arvey, Erik Larsson, Chris Sander, Christina S Leslie, and Debora S Marks. Target mRNA abundance dilutes microRNA and siRNA activity. Molecular Systems Biology, 6, April 2010. ISSN 1744-4292. doi: 10.1038/msb.2010.24.

15. Rémy Denzler, Vikram Agarwal, Joanna Stefano, David P. Bartel, and Markus Stoffel. Assessing the ceRNA Hypothesis with Quantitative Measurements of miRNA and Target Abundance. Molecular Cell, 54(5):766–776, June 2014. ISSN 10972765. doi: 10.1016/j.molcel.2014.03.045.

16. U. Ala, F. A. Karreth, C. Bosia, A. Pagnani, R. Taulli, V. Leopold, Y. Tay, P. Provero, R. Zecchina, and P. P. Pandolfi. Integrated transcriptional and competitive endogenous RNA networks are cross-regulated in permissive molecular environments. Proceedings of the National Academy of Sciences, 110(18):7154–7159, April 2013. ISSN 0027-8424, 1091-6490. doi: 10.1073/pnas.1222509110.

17. Marcella Cesana and George Q. Daley. Deciphering the rules of ceRNA networks. Proceedings of the National Academy of Sciences of the United States of America, 110(18): 7112–7113, April 2013. ISSN 1091-6490. doi: 10.1073/pnas.1305322110.

18. Rémy Denzler, Sean E. McGeary, Alexandra C. Title, Vikram Agarwal, David P. Bartel, and Markus Stoffel. Impact of MicroRNA Levels, Target-Site Complementarity, and Cooperativity on Competing Endogenous RNA-Regulated Gene Expression. Molecular Cell, 64(3):565–579, November 2016. ISSN 10972765. doi: 10.1016/j.molcel.2016.09.027.

19. Matteo Figliuzzi, Enzo Marinari, and Andrea De Martino. MicroRNAs as a Selective Channel of Communication between Competing RNAs: a Steady-State Theory. Biophysical Journal, 104(5):1203–1213, March 2013. ISSN 00063495. doi: 10.1016/j.bpj.2013.01.012.

20. Mor Nitzan, Avital Steiman-Shimony, Yael Altuvia, Ofer Biham, and Hanah Margalit. Interactions between Distant ceRNAs in Regulatory Networks. Biophysical Journal, 106(10): 2254–2266, May 2014. ISSN 00063495. doi: 10.1016/j.bpj.2014.03.040.

21. J. M. Robinson and W. A. Henderson. Modelling the structure of a ceRNA-theoretical, bipartite microRNA-mRNA interaction network regulating intestinal epithelial cellular pathways using R programming. BMC research notes, 11(1):19, January 2018. ISSN 1756-0500. doi: 10.1186/s13104-018-3126-y.

22. Markus List, Azim Dehghani Amirabad, Dennis Kostka, and Marcel H Schulz. Large-scale inference of competing endogenous rna networks with sparse partial correlation. Bioinformatics, 35(14):i596–i604, 2019.

23. Willian da Silveira, Ludivine Renaud, Jonathan Simpson, William Glen, Edward Hazard, Dongjun Chung, and Gary Hardiman. mirmapper: A tool for interpretation of mirna–mrna interaction networks. Genes, 9(9):458, 2018.

24. Xin Lai, Olaf Wolkenhauer, and Julio Vera. Understanding microRNA-mediated gene regulatory networks through mathematical modelling. Nucleic Acids Research, 44(13):6019–6035, July 2016. ISSN 0305-1048, 1362-4962. doi: 10.1093/nar/gkw550.

25. Leonardo Salmena, Laura Poliseno, Yvonne Tay, Lev Kats, and Pier Paolo Pandolfi. A ceRNA Hypothesis: The Rosetta Stone of a Hidden RNA Language? Cell, 146(3):353–358, August 2011. ISSN 00928674. doi: 10.1016/j.cell.2011.07.014.

26. Yvonne Tay, John Rinn, and Pier Paolo Pandolfi. The multilayered complexity of ceRNA crosstalk and competition. Nature, 505(7483):344–352, January 2014. ISSN 0028-0836, 1476-4687. doi: 10.1038/nature12986.

27. Ming Luo and Jun-Lin Guan. Focal adhesion kinase: a prominent determinant in breast cancer initiation, progression and metastasis. Cancer letters, 289(2):127–139, 2010.

28. Lisa Vettore, Rebecca L Westbrook, and Daniel A Tennant. New aspects of amino acid metabolism in cancer. British journal of cancer, pages 1–7, 2019.

29. Beatrice A Howard and Jacqueline M Veltmaat. Embryonic mammary gland development; a domain of fundamental research with high relevance for breast cancer research. Journal of Mammary Gland Biology and Neoplasia, 18(2):89–91, 2013.

30. Janet Piñero, Juan Manuel Ramírez-Anguita, Josep Saüch-Pitarch, Francesco Ronzano, Emilio Centeno, Ferran Sanz, and Laura I Furlong. The disgenet knowledge platform for disease genomics: 2019 update. Nucleic acids research, 48(D1):D845–D855, 2020.

31. Zhou Huang, Jiangcheng Shi, Yuanxu Gao, Chunmei Cui, Shan Zhang, Jianwei Li, Yuan Zhou, and Qinghua Cui. Hmdd v3. 0: a database for experimentally supported human microrna–disease associations. Nucleic acids research, 47(D1):D1013–D1017, 2018.

32. Gianluca Turcatel, Nicole Rubin, Ahmed El-Hashash, and David Warburton. Mir-99a and mir-99b modulate tgf-β induced epithelial to mesenchymal plasticity in normal murine mammary gland cells. PloS one, 7(1):e31032, 2012.

33. Su-Jin Yang, Su-Yu Yang, Dan-Dan Wang, Xiu Chen, Hong-Yu Shen, Xiao-Hui Zhang, Shan-Liang Zhong, Jin-Hai Tang, and Jian-Hua Zhao. The mir-30 family: versatile players in breast cancer. Tumor Biology, 39(3):1010428317692204, 2017.

34. Dingren Cao, Min Di, Jingjie Liang, Shuang Shi, Qiang Tan, and Zhengguang Wang. Microrna-183 in cancer progression. Journal of Cancer, 11(6):1315, 2020.

35. Da Wei Huang, Brad T Sherman, and Richard A Lempicki. Bioinformatics enrichment tools: paths toward the comprehensive functional analysis of large gene lists. Nucleic acids research, 37(1):1–13, 2009.

36. Brad T Sherman, Richard A Lempicki, et al. Systematic and integrative analysis of large gene lists using david bioinformatics resources. Nature protocols, 4(1):44, 2009.

37. Muhua Yang, Yuan Yao, Gabriel Eades, Yongshu Zhang, and Qun Zhou. Mir-28 regulates nrf2 expression through a keap1-independent mechanism. Breast cancer research and treatment, 129(3):983–991, 2011.

38. Daniela Schwarzenbacher, Christiane Klec, Barbara Pasculli, Stefanie Cerk, Beate Rinner, Michael Karbiener, Cristina Ivan, Raffaela Barbano, Hui Ling, Annika Wulf-Goldenberg, et al. Mir-1287-5p inhibits triple negative breast cancer growth by interaction with phosphoinositide 3-kinase cb, thereby sensitizing cells for pi3kinase inhibitors. Breast Cancer Research, 21(1):20, 2019.

39. Antonio Daniel Martinez-Gutierrez, David Cantú de León, Oliver Millan-Catalan, Jossimar Coronel-Hernandez, Alma D Campos-Parra, Fany Porras-Reyes, Angelica Exayana-Alderete, César López-Camarillo, Nadia J Jacobo-Herrera, Rosalio Ramos-Payan, et al. Identification of mirna master regulators in breast cancer. Cells, 9(7):1610, 2020.

40. Vasily N Aushev, Davide Degli Esposti, Eunjee Lee, Hector Vargas, Zdenko Herceg, Jun Zhu, and Jia Chen. mir-500a is involved in breast cancer-related gene expression pathways and associated with patients survival, 2016.

41. Katharina Uhr, Wendy JC Prager-van der Smissen, Anouk AJ Heine, Bahar Ozturk, Marijn TM van Jaarsveld, Antonius WM Boersma, Agnes Jager, Erik AC Wiemer, Marcel Smid, John A Foekens, et al. Micrornas as possible indicators of drug sensitivity in breast cancer cell lines. PloS one, 14(5), 2019.

42. Lei Huang, Xiaoqiao Tang, Xianbiao Shi, and Lei Su. mir-532-5p promotes breast cancer proliferation and migration by targeting rerg. Experimental and Therapeutic Medicine, 19 (1):400–408, 2020.

43. Chonggao Yin, Qingjie Mou, Xinting Pan, Guoxin Zhang, Hongli Li, and Yunbo Sun. Mir-577 suppresses epithelial-mesenchymal transition and metastasis of breast cancer by targeting rab25. Thoracic cancer, 9(4):472–479, 2018.

44. Chih-Hung Chou, Sirjana Shrestha, Chi-Dung Yang, Nai-Wen Chang, Yu-Ling Lin, Kuang-Wen Liao, Wei-Chi Huang, Ting-Hsuan Sun, Siang-Jyun Tu, Wei-Hsiang Lee, et al. mirtarbase update 2018: a resource for experimentally validated microrna-target interactions. Nucleic acids research, 46(D1):D296–D302, 2017.

45. Stefan Rüegger and Helge Großhans. MicroRNA turnover: when, how, and why. Trends in Biochemical Sciences, 37(10):436–446, October 2012. ISSN 09680004. doi: 10.1016/j.tibs.2012.07.002.

46. Ziynet Nesibe Kesimoglu and Serdar Bozdag. Inferring competing endogenous rna (cerna) interactions in cancer. bioRxiv, 2020.

47. Tayier Tuersong, Linlin Li, Zumureti Abulaiti, and Shumei Feng. Comprehensive analysis of the aberrantly expressed lncrna-associated cerna network in breast cancer. Molecular Medicine Reports, 19(6):4697–4710, 2019.

48. Xue Wang, Chundi Gao, Fubin Feng, Jing Zhuang, Lijuan Liu, Huayao Li, Cun Liu, Jibiao Wu, Xia Zheng, Xia Ding, et al. Construction and analysis of competing endogenous rna networks for breast cancer based on tcga dataset. BioMed Research International, 2020, 2020.

49. Jun Gong, Hui Jiang, Chang Shu, Mei-qin Hu, Yan Huang, Qin Liu, Rong-feng Li, and Yin-zhi Wei. Integrated analysis of circular rna-associated cerna network in cervical cancer: Observational study. Medicine, 98(34), 2019.

50. Weijun Liu and Xiaowei Wang. Prediction of functional microrna targets by integrative modeling of microrna binding and target expression data. Genome biology, 20(1):18, 2019.

51. Selcen Ari Yuka and Alper Yilmaz. A new network-based tool to analyse competing endogenous rnas. In Zhipeng Cai, Ion Mandoiu, Giri Narasimhan, Pavel Skums, and Xuan Guo, editors, Bioinformatics Research and Applications, pages 274–281, Cham, 2020. Springer International Publishing. ISBN 978-3-030-57821-3.

52. Stanislas Werfel, Simon Leierseder, Benjamin Ruprecht, Bernhard Kuster, and Stefan Engelhardt. Preferential microRNA targeting revealed by in vivo competitive binding and differential Argonaute immunoprecipitation. Nucleic Acids Research, 45(17):10218–10228, September 2017. ISSN 0305-1048, 1362-4962. doi: 10.1093/nar/gkx640.

53. Marcel Schulz Markus List. SPONGE, 2017.

54. NV Trang, Marc Choisy, T Nakagomi, NTM Chinh, YH Doan, T Yamashiro, JE Bryant, O Nakagomi, and DD Anh. Determination of cut-off cycle threshold values in routine rt– pcr assays to assist differential diagnosis of norovirus in children hospitalized for acute gastroenteritis. Epidemiology & Infection, 143(15):3292–3299, 2015.

