## Supplementary Figures and Tables for "Network based multifactorial modelling of miRNA-target interactions"

Selcen Ari Alper Yilmaz

2020-11-13

**4. Access to code** **13**

**REFERENCES** **14**

#### 2. Supplementary Figures

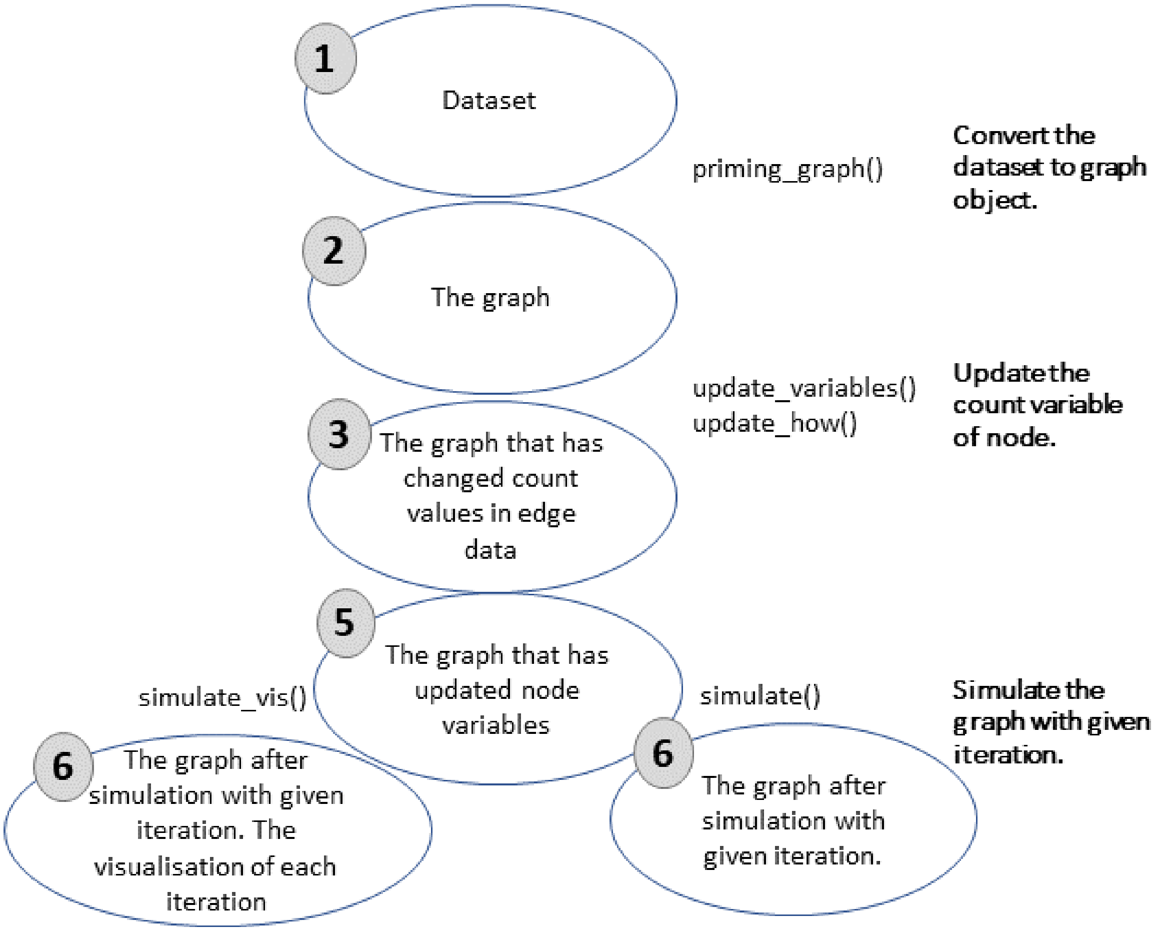

Figure S1: Workflow for simulation of competing endogenous RNA regulations. Graph object in steps 2-6 is saved and updated continuously.

##### 2.1 Sample dataset analysis in absence of interaction factors.

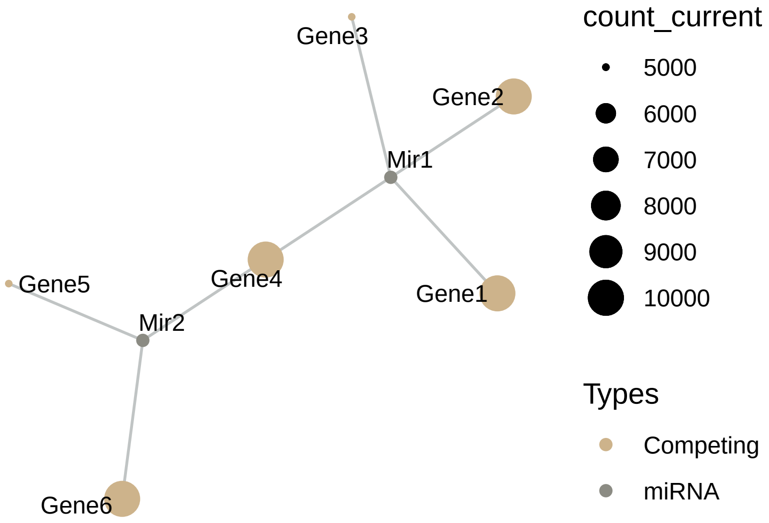

Figure S2: Sample Dataset in Steady-state. Initial expression levels in minsamp network (*Sample* network in manuscript). The network contains two miRNAs and 6 genes with arbitrarily chosen expression values. Refer to Table S1 for exact expression values

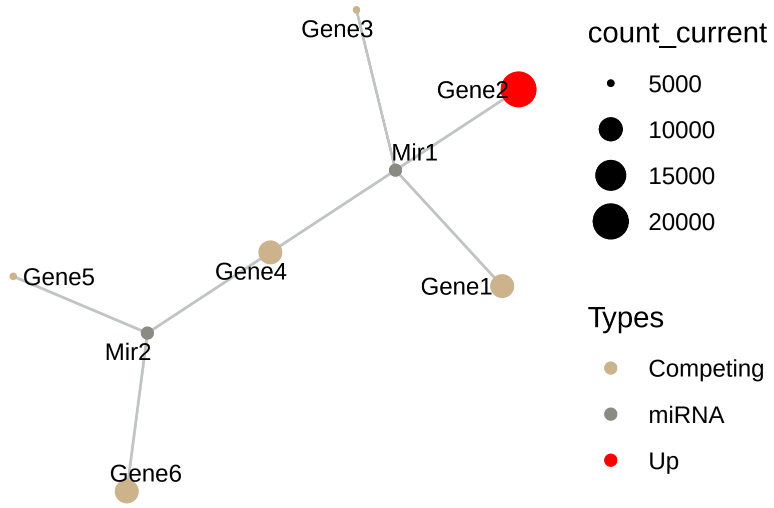

Figure S3: Gene2 Upregulation on Sample Dataset. Expression level of Gene2 is increased from 10,000 to 20,000 in order to demonstrate the calculation steps.

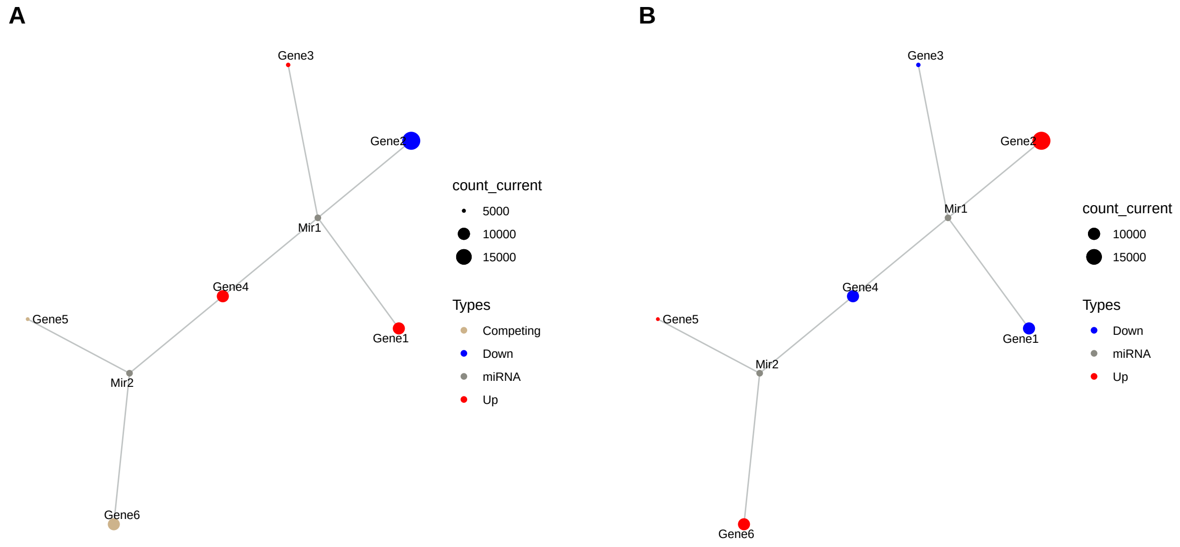

Figure S4: Sequential iteration of Sample data after up-regulation of Gene2. A) First response of system to Gene2 upregulation (2nd iteration). B) Spreading of perturbation on system (3th iteration)

#### 2.2 Calculations with interaction factors

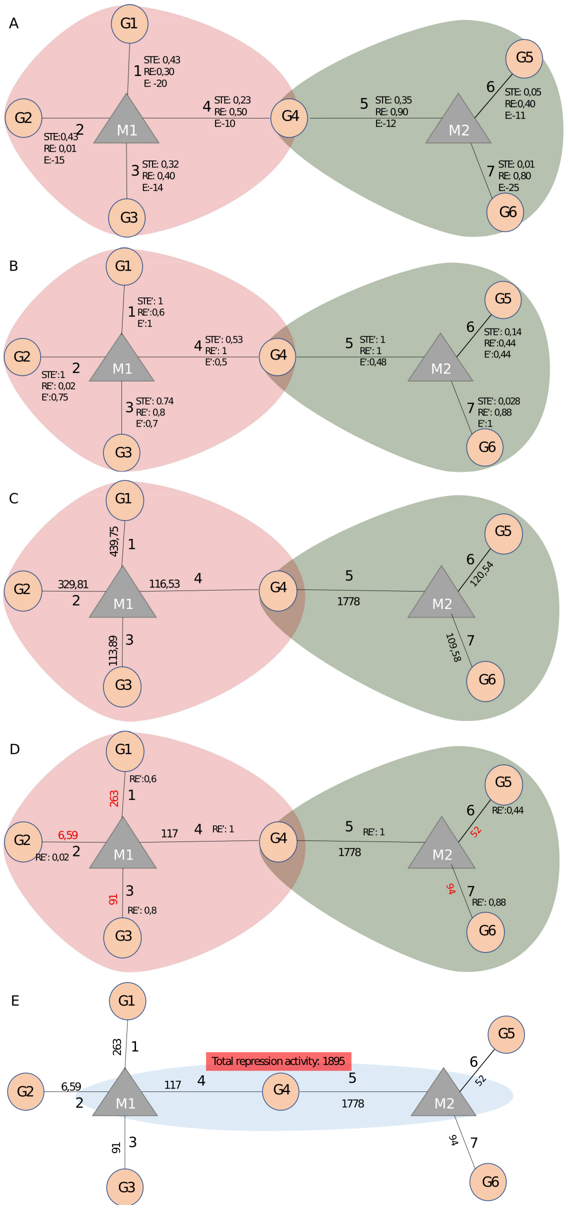

Figure S5: Calculation of initial miRNA repression activity using interaction parameters in Sample+ network. A) Interaction paramaters between genes and miRNAs in Sample+ are shown on network while expression levels can be found in Table S1. B) Interaction paramaters were updated after normalisation C) Amount of miRNA distributed to each mRNA according to mRNA levels and affinity parameters are shown on edges. D) Values on edges indicate repression activity. Red values indicate repression activity affected by region effect (RE) parameter. E) Total repression on G4 from two miRNAs is calculated by adding repression values originating from both miRNAs.

$G$, Gene; $M$, miRNA; $STE$, seed type effect; $RE$, Region Effect; $E$, Energy; $STE'$, normalized values of seed type efficiency coefficient; $RE'$, normalized values of region efficiency coefficient; $E'$, normalized values of energy coefficient. Numbering on edges match the pair order in Table S1.

##### 2.3 Sample+ dataset analysis with interaction factors.

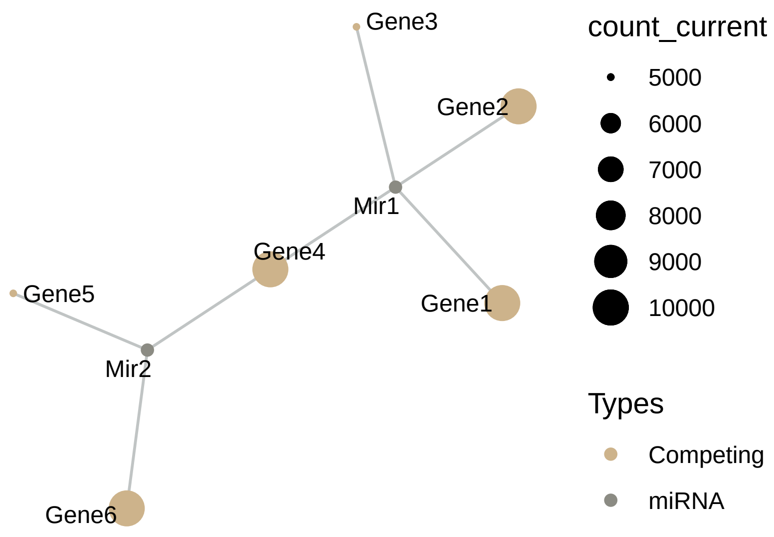

Figure S6: Sample+ in Steady-state. Interaction factors of Sample+ network are available in Table S1.

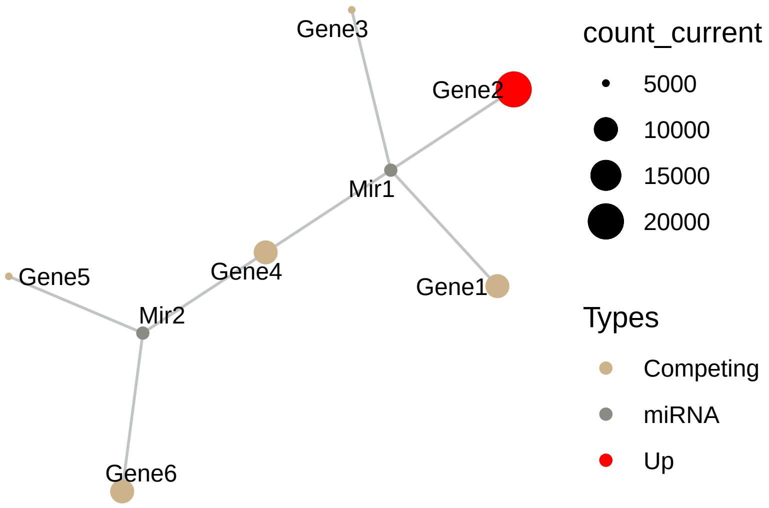

Figure S7: Perturbation in Sample+ network by two-fold increase in Gene2 expression level.

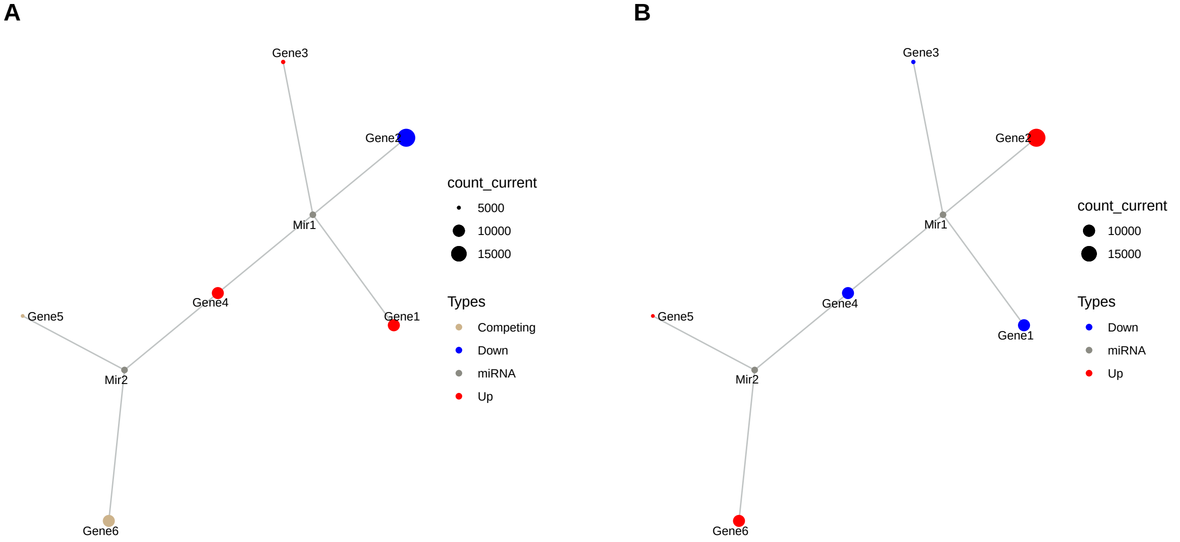

Figure S8: Sequential iteration of Sample+ A) First response of system to Gene2 upregulation (2nd iteration). B) Spreading of perturbation on system (3rd iteration). Although visualisation looks similar to Figure S4B, current counts of genes are drastically different.

#### 2.4 Common target perturbation in Sample+ dataset.

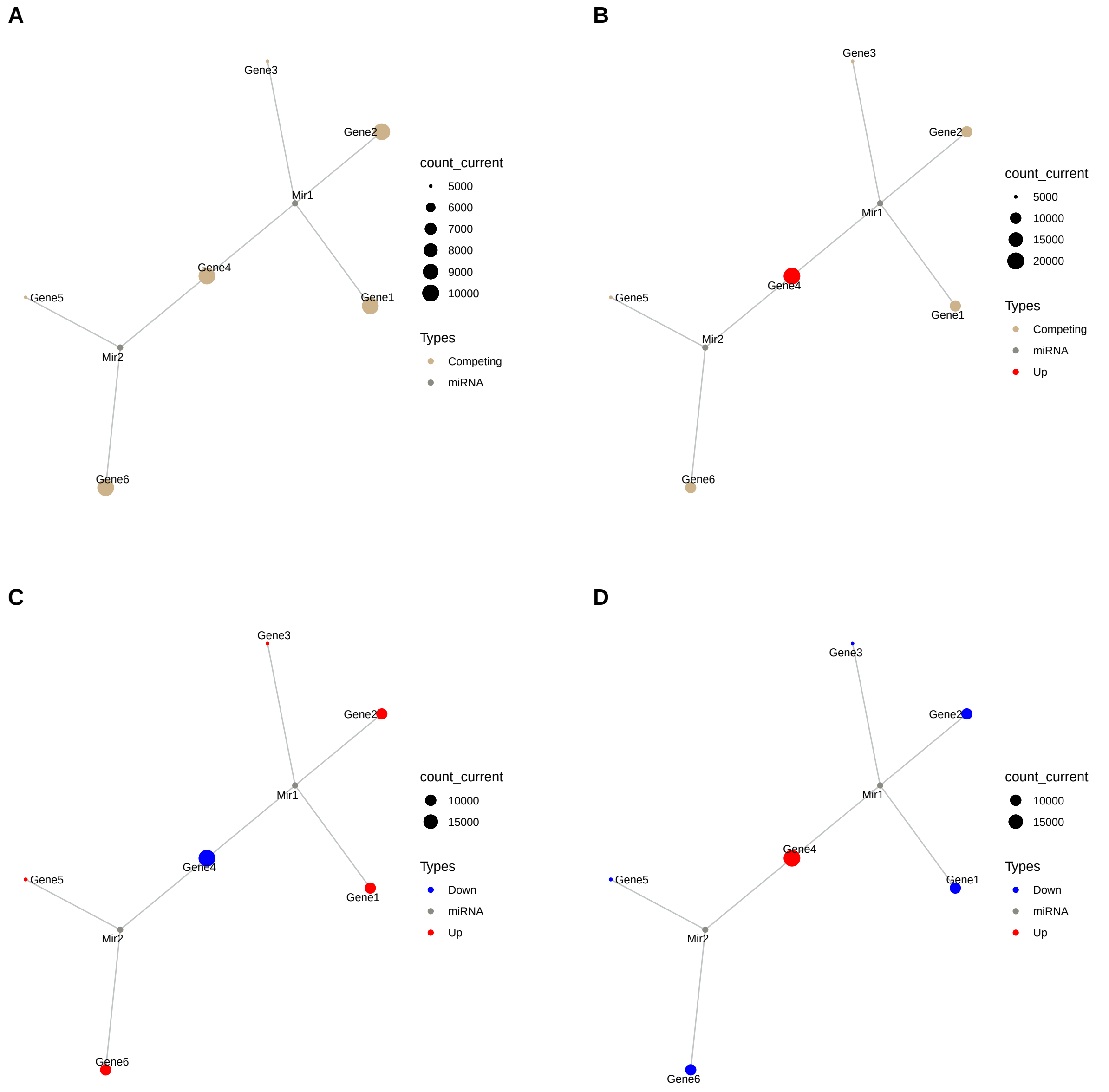

Figure S9: Perturbation of Gene4 and its effects on Sample+. A) Network at steady-state. B) Upregulation of Gene4. C) Primary response of network to upregulation of Gene4. D) Re-regulation of whole nodes on system (3th iteration)

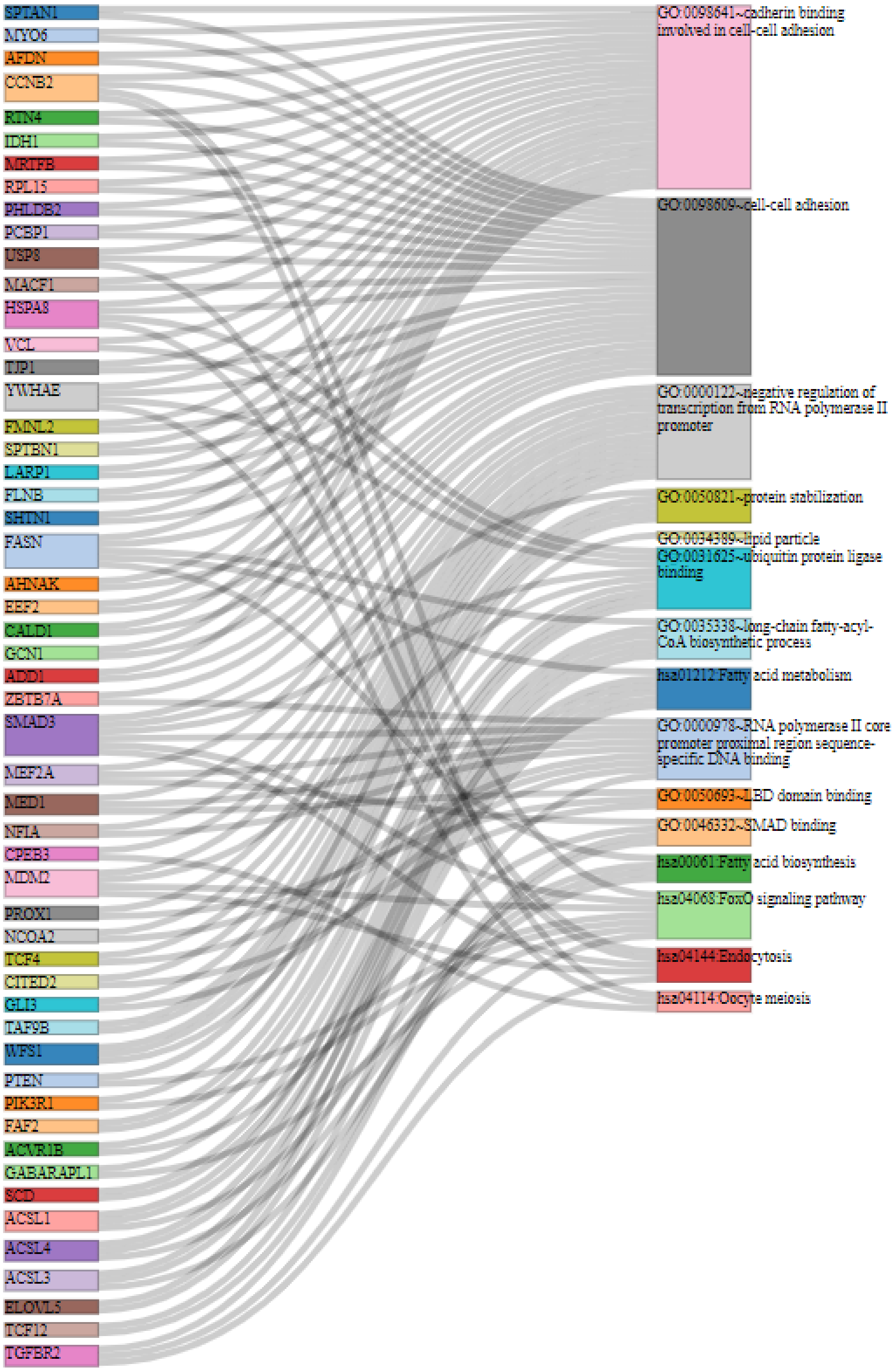

Figure S10: Sankey diagram represents top five KEGG and GO (molecular function and biological process) terms and genes enriched on these terms. Genes with single edge are not shown.

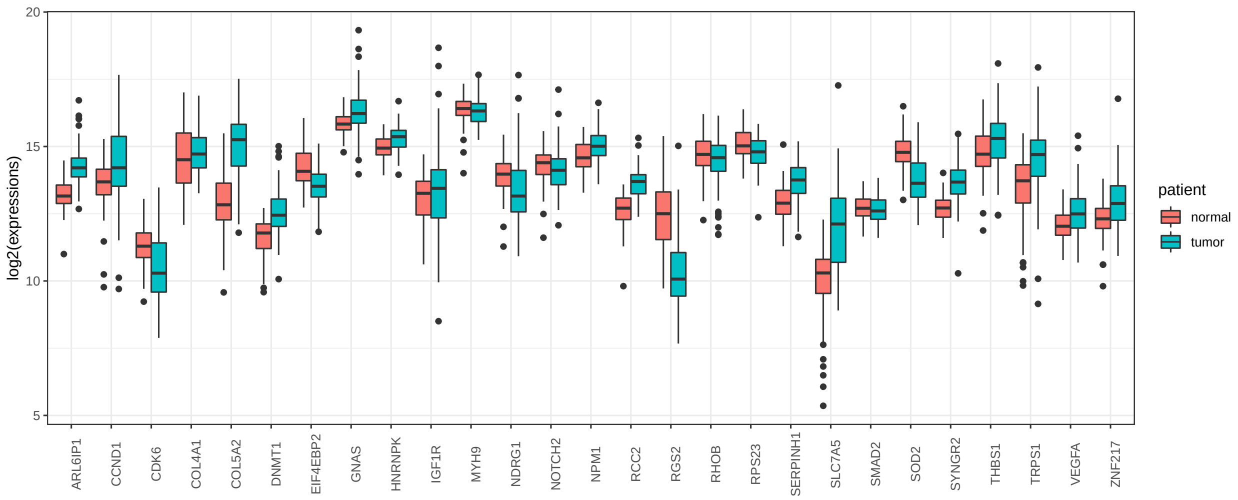

Figure S11: Log2 transformed expression levels of tumor-specific perturbing nodes in tumor and normal tissue samples of 87 patients.

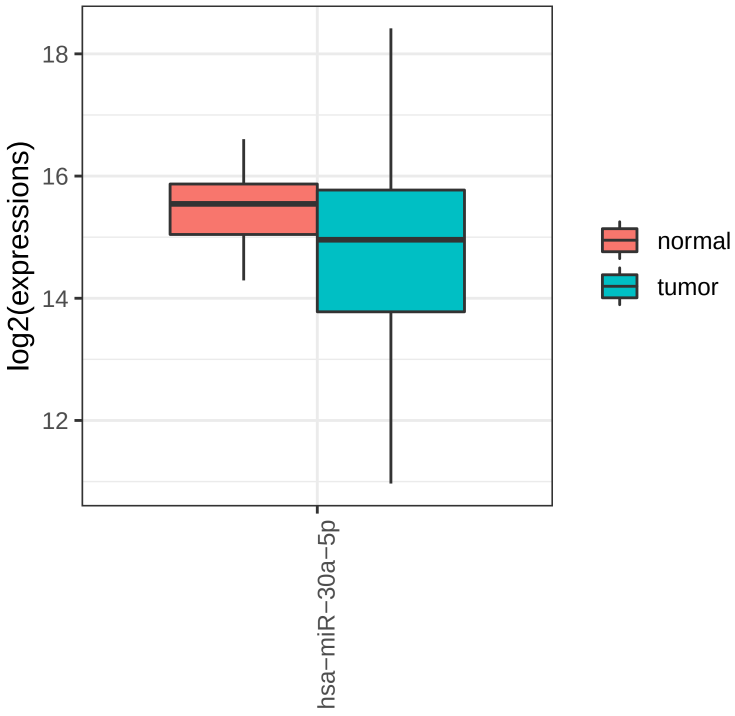

Figure S12: Log2 transformed expression level of miR-30a-5p in tumor and normal tissue samples of 87 patients.

### 3 Supplementary Tables

#### 3.1 *Sample+* dataset

Table S1: The parameters which affect miRNA:target interactions (i.e. seed type, region, energy) are provided in Sample+ dataset, while these factors are not utilized in simulation of Sample dataset.

| Competing | miRNA | Competing Expression | miRNA Expression | Seed Type Effect | Region Effect | Energy |
|---|---|---|---|---|---|---|
| `Gene1` | `Mir1` | 10000 | 1000 | 0.43 | 0.30 | -20 |
| `Gene2` | `Mir1` | 10000 | 1000 | 0.43 | 0.01 | -15 |
| `Gene3` | `Mir1` | 5000 | 1000 | 0.32 | 0.40 | -14 |
| `Gene4` | `Mir1` | 10000 | 1000 | 0.23 | 0.50 | -10 |
| `Gene4` | `Mir2` | 10000 | 2000 | 0.35 | 0.90 | -12 |
| `Gene5` | `Mir2` | 5000 | 2000 | 0.05 | 0.40 | -11 |
| `Gene6` | `Mir2` | 10000 | 2000 | 0.01 | 0.80 | -25 |

#### 3.2 Significant factors in miRNA:target interactions

Some of information about miRNA:target interactions were exhibited directly by high-throughput studies. On the other hand, we were examined other interacion factors based on different studies.

- (Helwak et al. 2013; Moore et al. 2015) reported the energy values in miRNA:target interactions.
- Comparisons of canonical seed types were evaluated by study of (Grimson et al. 2007), while functional and non-functional seed interactions were studied by (Bartel 2009) and (Betel et al. 2010).
- Numeric definition of target region location effect was performed based on studies of (Hausser et al. 2013) and (Helwak et al. 2013)

Table S2: Efficiency factors for seed types.

| seed type | seed type effect |
|---|---|
| 6-mer_noncanonical | 0.05 |
| 9-mer | 0.43 |
| 6-mer | 0.07 |
| 8-mer | 0.43 |
| 7-mer | 0.23 |
| none | 0.01 |
| 5-mer_noncanonical | 0.04 |
| 5-mer | 0.05 |
| 6-merA1_noncanonical | 0.05 |
| 7-mer-8m_noncanonical | 0.21 |
| 7-mer-8m | 0.25 |
| 8-mer_noncanonical | 0.35 |
| 7-merA1_noncanonical | 0.16 |
| 7-merA1 | 0.19 |
| 6-merA1 | 0.07 |

Table S3: Efficiency factors for binding regions on targets

| region | region effect |
|---|---|
| 3UTR | 0.84 |
| CDS | 0.42 |

| region | region effect |
|---|---|
| 3UTRCDS | 0.93 |
| 5UTR | 0.01 |
| 5UTRCDS | 0.42 |
| none | 0.01 |
| intron | 0.01 |
| CDS3UTR | 0.93 |
| CDS5UTR | 0.42 |
| exon_unclassified | 0.20 |
| CDS3UTRintron | 0.93 |
| 3UTRintron | 0.84 |
| CDSintron | 0.42 |
| 5UTRintron | 0.01 |
| 5UTR3UTR | 0.93 |
| CDS5UTR3UTR | 0.93 |

##### 3.3 Content of High-throughput experimental studies

Table S4: Context of miRNA:target pairs supported by High-throughput Experiments. CLEAR-CLiP and CLASH datasets were integrated as described in Section 2 of Supplementary Material and Method.

| Variable | Definition |
|---|---|
| cluster | Barcode from experimental method |
| chromosome | Chromosome of Target gene from raw data |
| start_position | Gene start position from raw data |
| end_position | Gene end position from raw data |
| strand | Gene strand |
| hgnc_symbol | Gene name (Symbol) |
| Ensembl_Gene_Id | Ensembl Gene Id of gene |
| Ensembl_Transcript_Id | Ensembl transcript id of mRNA of Target gene |
| target_seq | mRNA sequences targeted by miRNA |
| miRNA | miRNA id (from miRBase version 21 ) |
| miR_seq | miRNA sequence |
| seed_type | seed type of miRNA:target interaction |
| Energy | Energy of miRNA:target binding |
| HG38build_loc | Recent chromosomal location of Gene |
| Genome_build | Genome build of given chromosome, start and end positions |
| region | interaction location on target |
| region_effect | Coefficient of location on target |
| seed_type_effect | Coefficient for seed sequence of miRNA:target interaction |

##### 3.4 Variables of network object during simulation

As a result of simulation a dataset, a graph object is obtained that includes various variables in edge and node data. A graph object includes variables at Table S5.

Table S5: The context graph object during the process.

| Variables | Description |
|---|---|
| *Node Variables* | |
| name | node name |
| type | Competing or miRNA |
| node_id | in on graph object |
| initial_count | Initial Expression value of node |
| count_pre | Expression value of node at previous regulation |
| count_current | Existing expression value of node |
| changes_variable | Regulation of node (Up, down or steady) |
| *Edge Variables* | |
| Competing name | name of genes |
| miRNA name | name of miRNAs |
| Competing expression | Expression values of competing elements at steady-state |
| miRNA expression | Expression values of miRNA elements at steady-state |
| energy | coefficient of miRNA:target interactions (binding affinity) |
| seed type | coefficient of miRNA:target interactions (binding affinity) |
| region | coefficient of miRNA:target interactions (degradation efficiency) |
| afff factor | coefficient scaled and combined affinity factor |
| degg factor | coefficient scaled and combined degradation factor |
| comp_count_list | list of competing expression for each iteration |
| comp_count | pre: competing expression at previous iteration; current: competing expression at present iteration |
| mirna_count_list | list of miRNA expression for each iteration |
| mirna_count | pre: miRNA expression at previous iteration; current: miRNA expression at present iteration |
| effect | pre: total miRNA reppressive effect on individual target at previous iteration ; current: miRNA reppressive effect on individual target at present iteration |
| effect_list | list of miRNA reppressive effect on individual target for each iteration |

### 4. Access to code

All codes are available at github repo.

### REFERENCES

Bartel, David P. 2009. “MicroRNAs: Target Recognition and Regulatory Functions.” *Cell* 136 (2): 215–33. https://doi.org/10.1016/j.cell.2009.01.002.

Betel, Doron, Anjali Koppal, Phaedra Agius, Chris Sander, and Christina Leslie. 2010. “Comprehensive Modeling of microRNA Targets Predicts Functional Non-Conserved and Non-Canonical Sites.” *Genome Biology* 11 (8): R90.

Grimson, Andrew, Kyle Kai-How Farh, Wendy K. Johnston, Philip Garrett-Engele, Lee P. Lim, and David P. Bartel. 2007. “MicroRNA Targeting Specificity in Mammals: Determinants Beyond Seed Pairing.” *Molecular Cell* 27 (1): 91–105. https://doi.org/10.1016/j.molcel.2007.06.017.

Hausser, J., A. P. Syed, B. Bilen, and M. Zavolan. 2013. “Analysis of CDS-Located miRNA Target Sites Suggests That They Can Effectively Inhibit Translation.” *Genome Research* 23 (4): 604–15. https://doi.org/10.1101/gr.139758.112.

Helwak, Aleksandra, Grzegorz Kudla, Tatiana Dudnakova, and David Tollervey. 2013. “Mapping the Human miRNA Interactome by CLASH Reveals Frequent Noncanonical Binding.” *Cell* 153 (3): 654–65. https://doi.org/10.1016/j.cell.2013.03.043.

Moore, Michael J., Troels K. H. Scheel, Joseph M. Luna, Christopher Y. Park, John J. Fak, Eiko Nishiuchi, Charles M. Rice, and Robert B. Darnell. 2015. “miRNA-Target Chimeras Reveal miRNA 3’-End Pairing as a Major Determinant of Argonaute Target Specificity.” *Nature Communications* 6 (November): 8864. https://doi.org/10.1038/ncomms9864.
