## Supplementary methods for "Network based multifactorial modelling of miRNA-target interactions"

### Supplementary Materials And Methods

Selcen Ari Alper Yilmaz

2020-11-16

**3. Functions defined for ceRNA models and workflow of method** **9**

**4. Perturbation analysis of Real/Real+ networks** **10**

REFERENCES **10**

The functions that are used in the this study are developed with R and are available in Bioconductor.

#### 1. Determination of perturbation efficiencies of nodes in network.

`find_node_perturbation` function runs `calc_perturbation` function for each node with given parameters. In this particular example upregulation parameter is two, iteration parameter is three. By using formulas described in manuscript perturbation efficiency and number of affected nodes (perturbed count) are calculated. Results for Sample+ network are shown below.

```r
library(ceRNAnetsim)
data("minsamp")  # refers Sample and Sample+ data

sample_graph <- priming_graph(minsamp, competing_count = Competing_expression,
    miRNA_count = miRNA_expression, aff_factor = c(energy,
        seed_type), deg_factor = region)

find_node_perturbation(sample_graph, how = 2, cycle = 3,
    limit = 0.1) %>% as_tibble() %>% select(name, perturbation_efficiency,
    perturbed_count)
```

```
## # A tibble: 8 x 3
##   name  perturbation_efficiency perturbed_count
##   <chr>                   <dbl>           <dbl>
## 1 Gene1                  0.132                2
## 2 Gene2                  0.198                3
## 3 Gene3                  0.0555               2
## 4 Gene4                  0.197                4
## 5 Gene5                  0.143                1
## 6 Gene6                  0.131                1
## 7 Mir1                   0.806                3
## 8 Mir2                   2.80                 3
```

### 2. Additional data manipulation steps

#### 2.1 Arrangement of CLASH dataset

CLASH dataset was retrieved from Helwak et. al (Helwak et al. 2013).

```r
clash_url <- "https://www.ncbi.nlm.nih.gov/pmc/articles/PMC3650559/bin/mmc1.txt"

clashelwak <- read_tsv(clash_url, comment = "#", skip = 1)

# hg19
```

###### Query of Human Genome 19.

Human genome 19 information was handled through biomaRt package.

```r
library(biomaRt)  #can be installed via BiocManager

# listMarts(host = 'http://grch37.ensembl.org')
# #HG19
ensemblgrch37 <- useMart(host = "http://grch37.ensembl.org",
    biomart = "ENSEMBL_MART_ENSEMBL", dataset = "hsapiens_gene_ensembl")

hg19 <- getBM(attributes = c("ensembl_transcript_id",
    "ensembl_gene_id", "chromosome_name", "start_position",
    "end_position", "hgnc_symbol", "entrezgene_id",
    "strand"), mart = ensemblgrch37)
```

##### Adding miRNA and gene information

```
clashelwak <- clashelwak %>% separate(microRNA_name,
    c("Barcode", "Database", "mirna_name", "type"),
    sep = "_") %>% separate(mRNA_name, c("Ensembl_Gene_Id",
    "Ensembl_Transcript_Id", "Hugo_Symbol", "mRNA_Type"),
    sep = "_") %>% rename_with(~str_replace(., "'",
    "_"), contains("UTR"))
```

miRNA releases are obtained from miRBase. In this step, release 21 (in Human genome 38) was downloaded.

```
mirbase_url <- "ftp://mirbase.org/pub/mirbase/21/genomes/hsa.gff3"

mirbasehg38 <- read_tsv(mirbase_url, comment = "#",
    col_names = FALSE) %>% filter(X3 != "miRNA_primary_transcript") %>%
    separate(X9, c("ID", "Alias", "Name", "Precusor"),
        sep = ";") %>% mutate(ID = substr(ID, 4, length(ID)),
    Alias = substr(Alias, 7, length(Alias)), Name = substr(Name,
        6, length(Name)), Precusor = substr(Precusor,
        14, length(Precusor))) %>% dplyr::select(chr = X1,
    start = X4, end = X5, strand = X7, ID, Alias, Name,
    Precusor)
```

Since CLASH dataset is published according to Human Genome 19 version, we used miRBase release 15.

```
mirbase15_url <- "ftp://mirbase.org/pub/mirbase/15/database_files/mirna_mature.txt.gz"

clashelwakfinal <- read_tsv(mirbase15_url, col_names = FALSE) %>%
    filter(str_starts(X2, "hsa")) %>% dplyr::select(mirna_ID = X2,
    mirbase_ID = X3) %>% inner_join(mirbasehg38 %>%
    dplyr::select(ID, Name), by = c(mirbase_ID = "ID")) %>%
    dplyr::select(mirbase_ID, Name) %>% distinct() %>%
    inner_join(clashelwak, by = c(mirbase_ID = "Barcode")) %>%
    dplyr::select(Name, miRNA_seq, Ensembl_Gene_Id,
        Ensembl_Transcript_Id, Hugo_Symbol, mRNA_seq_extended,
        chimeras_decompressed, seed_type, seed_basepairs,
        folding_class, seq_ID, folding_energy, `5_UTR`,
        CDS, `3_UTR`) %>% inner_join(hg19, by = c(Ensembl_Gene_Id = "ensembl_gene_id",
    Ensembl_Transcript_Id = "ensembl_transcript_id",
    Hugo_Symbol = "hgnc_symbol")) %>% mutate(region1 = ifelse(`5_UTR` ==
    "1", "5UTR", " "), region2 = ifelse(`3_UTR` ==
    "1", "3UTR", " "), region3 = ifelse(CDS == "1",
    "CDS", " ")) %>% unite(region, c(region1, region2,
    region3), sep = "||") %>% dplyr::select(chromosome_name,
    start_position, end_position, strand, Hugo_Symbol,
    Ensembl_Gene_Id, Ensembl_Transcript_Id, mRNA_seq_extended,
    Name, miRNA_seq, seq_ID, seed_type, seed_basepairs,
    folding_class, folding_energy, region)
```

##### Converting CLASH data to human genome 38 build.

There are different liftover methods for conversion among Human Genome builds. We preffered to use UCSC liftover tool

```r
# Obtaining chromosomal locations from miRNA:target
# interaction dataset.

clashelwakfinal %>% dplyr::select(1, 2, 3) %>% unite(start_end,
    c("start_position", "end_position"), sep = "-") %>%
    mutate(Chromosome = paste0("chr", chromosome_name,
        "")) %>% unite(chromosome_name, c("Chromosome",
    "start_end"), sep = ":") %>% write_tsv("lift19.txt",
    col_names = FALSE)

# In liftover site, Original Assembly was chosen as
# hg19 and New Assembly was chosen as hg38. After
# we uploaded 'lift19.txt' file to UCSC Liftover
# site, the output 'View Conversions' was
# downloaded as hg38clashcomp (new locations of the
# genes on HG38), 'Display failure file' was
# download as 'lift19_del.txt' (deleted regions on
# the HG38 genome build)

lift19_del <- read_tsv("data/lift19_del.txt", col_names = FALSE,
    comment = "#") %>% rename(chromosome_loc = X1) %>%
    dplyr::filter(str_starts(chromosome_loc, "chr")) %>%
    separate(chromosome_loc, c("Chr", "End"), "-",
        remove = TRUE) %>% separate(Chr, c("Chr", "Start"),
    ":", remove = TRUE) %>% mutate(Start = as.numeric(Start),
    End = as.numeric(End))

# removing deleted location from CLASH dataset

clashelwakfinal <- clashelwakfinal %>% mutate(Chromosome = paste0("chr",
    chromosome_name, "")) %>% dplyr::anti_join(lift19_del,
    by = c(Chromosome = "Chr", start_position = "Start",
        end_position = "End"))
# adding new location information:

clashelwakfinal <- clashelwakfinal %>% bind_cols(read_tsv("data/hg38clashcomp.txt",
    col_names = FALSE))

# Arrangement in dataset

clashelwakfinal <- clashelwakfinal %>% rename(HG38build_loc = X1) %>%
    mutate(Genom_build = "hg19") %>% dplyr::select(cluster = seq_ID,
    chromosome = Chromosome, start_position, end_position,
    strand, hgnc_symbol = Hugo_Symbol, Ensembl_Gene_Id,
    Ensembl_Transcript_Id, target_seq = mRNA_seq_extended,
    miRNA = Name, miR_seq = miRNA_seq, seed_type, seed_type2 = seed_basepairs,
    seed_type3 = folding_class, Energy = folding_energy,
    HG38build_loc, Genom_build, region) %>% mutate(strand = as.character(strand))
```

###### Interpreting the CLASH seed structures in dataset

```
clashelwakfinal <- clashelwakfinal %>% mutate(seed_type = ifelse(seed_type ==
    "noncanonical_seed" & seed_type2 > 4 & seed_type3 ==
    "I", paste0(seed_type2, "-mer"), seed_type), seed_type = ifelse(seed_type ==
    "noncanonical_seed" & seed_type2 > 4 & seed_type3 ==
    "II", paste0(seed_type2, "-mer_noncanonical"),
    seed_type), seed_type = ifelse(seed_type == "noncanonical_seed" &
    seed_type2 > 4 & seed_type3 == "III", paste0(seed_type2,
    "-mer_noncanonical"), seed_type), seed_type = ifelse(seed_type ==
    "noncanonical_seed" & seed_type2 > 4 & seed_type3 ==
    "IV", paste0(seed_type2, "-mer_noncanonical"),
    seed_type), seed_type = ifelse(startsWith(seed_type,
    "no"), "none", seed_type)) %>% dplyr::select(-seed_type2,
    -seed_type3)
```

#### 2.2 Arrangement of CLEAR-CLiP Dataset (Moore et al. 2015)

CLEAR-CLiP dataset was retrieved from Moore et.al. Please note that we need Supplementary Data 4, however there is error in website, you need to click on Supplementary Data 3 to download the data.

```
clear_clip_url <- "https://static-content.springer.com/esm/art%3A10.1038%2Fncomms9864/MediaObjects/4146

download.file(url = clear_clip_url, destfile = "CLEAR-CLIP.xlsx",
    mode = "wb")

clearclip <- readxl::read_excel("CLEAR-CLIP.xlsx",
    1, skip = 23)

# Clearclip hg18
```

###### Query of Human Genome 18

```
# HG18
library(biomaRt)

# listMarts(host = 'may2009.archive.ensembl.org')
ensembl54 = useEnsembl(host = "may2009.archive.ensembl.org",
    biomart = "ensembl", dataset = "hsapiens_gene_ensembl",
    mirror = "asia")

hg18 <- getBM(attributes = c("ensembl_transcript_id",
    "ensembl_gene_id", "chromosome_name", "start_position",
    "end_position", "hgnc_symbol", "entrezgene_id",
    "strand"), mart = ensembl54)

# should there be any connection problems, read the
# pre-saved version of hg18 gene information by
```

```
# commenting out the following line

# hg18 <- read_tsv('data/hg18.txt') %>%
# rename_with(~ tolower(.x) %>% str_replace_all('
# ','_') %>% str_remove('_\\(bp\\)')) %>%
# rename(start_position=gene_start,
# end_position=gene_end) %>%
# filter(!is.na(entrezgene_id)) %>%
# filter(!is.na(hgnc_symbol))
```

##### Adding Genome Information to dataset

```
clearclipfinal <- hg18 %>% inner_join(clearclip, by = c(entrezgene_id = "gene.id",
    hgnc_symbol = "gene.symbol")) %>% distinct()
```

##### Converting human genome build

```
# Obtaining chromosomal locations from miRNA:target
# interaction dataset.

lift18 <- clearclipfinal %>% unite(start_end, c("start_position",
    "end_position"), sep = "-") %>% unite(location,
    c("chr", "start_end"), sep = ":") %>% dplyr::select(location)

# you can save current state of lift18 data frame
# write_tsv(lift18, 'data/lift18.txt')
```

Please note that we have provided the hg18 liftover dataset under the data folder (lift18), in case of server errors/changes may occur. If you had problems generating lift18 data frame, please run the following code.

```
lift18 <- read_tsv("data/lift18.txt")
```

```
# In liftover site, Original Assembly was chosen as
# hg18 and New Assembly was chosen as hg38. After
# we uploaded 'lift18.txt' file to UCSC Liftover
# site, the output 'View Conversions' was
# downloaded as 'hg38clearclip.txt' (new locations
# of the genes on HG38), 'Display failure file' was
# download as 'lift18_del.txt' (deleted regions on
# the HG38 genome build)

deleted_lift18 <- read_tsv("data/lift18_del.txt", comment = "#",
    col_names = FALSE) %>% rename(Chromosome_loc = X1) %>%
    dplyr::filter(str_starts(Chromosome_loc, "chr")) %>%
    separate(Chromosome_loc, c("Chr", "End"), "-",
        remove = TRUE) %>% separate(Chr, c("Chr", "Start"),
    ":", remove = TRUE) %>% mutate(Start = as.numeric(Start),
    End = as.numeric(End))
```

```r
# removing deleted location from CLEAR-CLiP dataset

clearclipfinal <- clearclipfinal %>% dplyr::anti_join(deleted_lift18,
    by = c(chr = "Chr", start_position = "Start", end_position = "End"))

hg38clearclip <- read_tsv("data/hg38clearclip.txt",
    col_names = FALSE)

# adding new location information:
clearclipfinal <- clearclipfinal %>% bind_cols(hg38clearclip) %>%
    rename(HG38build_loc = X1) %>% mutate(Genom_build = "hg18") %>%
    dplyr::select(cluster = cluster.ID, chromosome = chr,
        start_position, end_position, strand = strand.y,
        hgnc_symbol, Ensembl_Gene_Id = ensembl_gene_id,
        Ensembl_Transcript_Id = ensembl_transcript_id,
        target_seq = target.map, miRNA, miR_seq = miR.map,
        seed_type = "seed match", Energy = MFE, HG38build_loc,
        Genom_build, region)
```

###### Seed type manipulation in CLEAR-CLiP dataset

In CLEAR-CLiP dataset, seed types were shown in detail. We adjusted as canonical and non-canonical.

```r
clipdata_seed <- data_frame(seed_type = c("5mer_1",
    "5mer_2", "5mer_3", "6mer", "6mer.indel", "6mer.mm",
    "6mer_off.mm", "6merA1", "6merA1.indel", "6merA1.mm",
    "7merA1", "7merA1.indel", "7merA1.mm", "7merm8",
    "7merm8.indel", "7merm8,mm", "8mer", "8mer.indel",
    "8mer.mm", "NA"), seed_type_com = c("5-mer", "5-mer_noncanonical",
    "5-mer_noncanonical", "6-mer", "6-mer_noncanonical",
    "6-mer_noncanonical", "6-mer_noncanonical", "6-merA1",
    "6-merA1_noncanonical", "6-merA1_noncanonical",
    "7-merA1", "7-merA1_noncanonical", "7-merA1_noncanonical",
    "7-mer-8m", "7-mer-8m_noncanonical", "7-mer-8m_noncanonical",
    "8-mer", "8-mer_noncanonical", "8-mer_noncanonical",
    "none"))

clearclipfinal <- clearclipfinal %>% inner_join(clipdata_seed,
    by = "seed_type") %>% dplyr::select(1:11, seed_type = seed_type_com,
    Energy, HG38build_loc, Genom_build, region)
```

##### 2.3 Integration of two experimental dataset

CLASH and CLEAR-CLiP data were processed heavily and resulted in similar format after which they were concatenated.

```r
experimentalmirnagene <- bind_rows(clashelwakfinal,
    clearclipfinal) %>% distinct()
```

#### 2.4 Adding Coefficients of Interaction factors

Energy values in miRNA:target pairs are represented by high-throughput studies (Helwak et al. 2013; Moore et al. 2015) which are utilized in this study. On the other hand, we have specified the other interaction factors, seed type and location of binding region on the target, as numeric values based on the previous studies.(Grimson et al. 2007) have compared the seed types' effect on target repression with few miRNA had canonical seed pairing in their study. Additionally, (Bartel 2009) and (Betel et al. 2010) have studied on functional and non-functional seed interactions. Based on results of these studies we have arranged seed types of miRNA:target interactions as numeric values. We also have redefined location of binding region on the target as numeric values, based on studies of (Hausser et al. 2013) and (Helwak et al. 2013). With this process, we have handled this integrated dataset in context of competitor behaviors and functionality of interactions.

In this step we added numeric interaction values as following

Firstly, we organized these values due to the fact that the regions were defined differently in two dataset. After that, region effect was added as numeric values (shown in Table S3).

```
experimentalmirnagene <- experimentalmirnagene %>%
    mutate(region2 = str_replace_all(region, "NA",
        ""), region3 = str_replace_all(region2, "\\|",
        ""), region = str_replace_all(region3, c(`3'UTR` = "3UTR",
        `5'UTR` = "5UTR"))) %>% dplyr::select(-region2,
    -region3) %>% mutate(region_effect = case_when(region %in%
    c("3UTRCDS", "CDS3UTR", "5UTR3UTR", "CDS5UTR3UTR",
        "CDS3UTRintron") ~ "0.93", region %in% c("5UTRCDS",
    "CDS5UTR") ~ "0.42", region %in% c("CDS", "CDSintron") ~
    "0.42", region %in% c("3UTR", "3UTRintron") ~ "0.84",
    region %in% c("5UTR", "5UTRintron") ~ "0.01", region %in%
        c("intron", "") ~ "0.01", region %in% c("exon_unclassified",
        "") ~ "0.2", TRUE ~ NA_character_)) %>% mutate(region_effect = as.double(region_effect))
```

Secondly, we organized seed type interactions in *Seed type manipulation* section for CLEAR-CLiP dataset to show as found in CLASH dataset. Same type formatted values added dataset as numeric values (shown in Table S2).

```
seed_type_effect <- data_frame(seed_type = c("5-mer",
    "5-mer_noncanonical", "6-mer", "6-mer_noncanonical",
    "6-merA1", "6-merA1_noncanonical", "7-mer", "7-mer_noncanonical",
    "7-merA1", "7-merA1_noncanonical", "7-mer-8m",
    "7-mer-8m_noncanonical", "8-mer", "8-mer_noncanonical",
    "9-mer", "9-mer_noncanonical", "none"), seed_type_effect = c(0.05,
    0.04, 0.07, 0.05, 0.07, 0.05, 0.23, 0.19, 0.19,
    0.16, 0.25, 0.21, 0.43, 0.35, 0.43, 0.35, 0.01))
experimentalmirnagene <- experimentalmirnagene %>%
    inner_join(seed_type_effect, by = "seed_type")
```

The methods about miRNA:target interactions are based a basic principle that is reading after isolation of miRNA:target chimerics. The datasets contain all the chimeric miRNA:target structures found in the medium during the experiment. On the other hand, it could be said that the reading is performed as snapshot.Because of that, the methods can provide different chimeric interactions the same miRNA:target pair. We have preferred to select most effective interaction parameters for the same miRNA:target pairs that can exhibit various interactions. The step is performed as:

```r
experimentalmirnagene <- experimentalmirnagene %>%
    dplyr::select(miRNA, Ensembl_Gene_Id, hgnc_symbol,
        Energy, seed_type_effect, region_effect) %>%
    distinct() %>% group_by(Ensembl_Gene_Id, miRNA) %>%
    mutate(seed_type_effect = ifelse(seed_type_effect ==
        max(seed_type_effect), seed_type_effect, max(seed_type_effect)),
        Energy = ifelse(Energy == min(Energy), Energy,
            min(Energy)), region_effect = ifelse(region_effect ==
            max(region_effect), region_effect, max(region_effect))) %>%
    distinct()
```

```r
# the data frame generated up to now should match
# the data frame used in manuscript so the output
# should be 'zero' different rows.
setdiff(experimentalmirnagene, readRDS("data/experimentalmirnagene.RDS"))
```

The context of dataset is shown in Table S5 in Supplementary Tables.

### 3. Functions defined for ceRNA models and workflow of method

We defined the functions that can be used with R programming. Briefly, these functions process a given miRNA:gene dataset and convert to graph object. All values that are significant in miRNA:target interactions are stored in edge variables and processed with formulations that are given in previous section. The functions and steps of approach are explained as following (Figure S1) :

**Conversion of dataset**: `priming_graph()` function processes the given dataset that includes competing elements in first variable and repressive element in second variable. If the affinity and/or degradation factors are specified in the function, factors are taken into account, are processed with defaults in vice versa. The formulations that are given in equations (1-4) are performed in this function. This step gives the graph object which contains efficiency values of miRNA:competing target pairs in steady-state in terms of amount. It is assumed that the initial target amounts in the dataset is observed after the repressive activity of miRNAs in steady-state.

**Transition of variables in graph**: In the previous step, the calculations are performed in the edge variables of the graph object. However, the graph object allows to use node variables, while the node features are handled to the graph. In this direction, update_nodes function carries the amount values to node variables This step must be applied with "once" option because it is primary process.

**Trigger change in graph**: The dataset are assumed as steady-state in previous step and the efficiency coefficients are calculated according to this acceptance. In the network that is found in steady-state conditions, the change is applied to the graph object for perturbation of steady-state. To provide the perturbation in the network the work-flow offer two methods: *update_variables* and *update_how*. The first, a new dataset that is contained competing and repressive element names and current values of these can be processed with update_variables. The second option, the amount of the given node name in update_how function can be changed according to "how" argument.

**Simulation of competing behavior of targets**: After the change in the steady-state conditions, the network elements try to gain steady-state again. This process progresses as repeating of regulations after the spreading the changes in the network. In this step, simulation of regulations according to given cycle count in simulate function is applied. After each simulation cycle, the miRNA repression values are re-calculated and the current values of competing elements are found and saved. The process is performed in the edge data and at the same time outputs of the calculations are carried from edge to node data.

The node elements in the dataset are handled as two type; repressive (miRNAs) and competing (targets). It is assumed in approach that while targets are degrading or inhibiting by miRNAs continuously, miRNAs reversibly used. If the trigger of the network is a miRNA, it maintains the current value of amount that provides by user. On the contrary, it tries to help this process to provide steady-state through the regulations on its amount, if a competing element is used as a trigger. The functions that are used in the this study are developed with R and are available in Bioconductor.

### 4. Perturbation analysis of Real/Real+ networks

To evaluate significant nodes through parallel processing in breast cancer patient network, perturbation on all nodes were triggered by 3-fold up-regulation with 10 iteration (cycle) due to small diameter of Real and Real+ networks. We analyzed perturbation efficiencies of all nodes in situation that accepts nodes with more than one percent change in expression as re-regulated (i.e, limit=1 argument in *find_node_perturbation()* function).

### REFERENCES

Bartel, David P. 2009. "MicroRNAs: Target Recognition and Regulatory Functions." *Cell* 136 (2): 215–33. https://doi.org/10.1016/j.cell.2009.01.002.

Betel, Doron, Anjali Koppal, Phaedra Agius, Chris Sander, and Christina Leslie. 2010. "Comprehensive Modeling of microRNA Targets Predicts Functional Non-Conserved and Non-Canonical Sites." *Genome Biology* 11 (8): R90.

Grimson, Andrew, Kyle Kai-How Farh, Wendy K. Johnston, Philip Garrett-Engele, Lee P. Lim, and David P. Bartel. 2007. "MicroRNA Targeting Specificity in Mammals: Determinants Beyond Seed Pairing." *Molecular Cell* 27 (1): 91–105. https://doi.org/10.1016/j.molcel.2007.06.017.

Hausser, J., A. P. Syed, B. Bilen, and M. Zavolan. 2013. "Analysis of CDS-Located miRNA Target Sites Suggests That They Can Effectively Inhibit Translation." *Genome Research* 23 (4): 604–15. https://doi.org/10.1101/gr.139758.112.

Helwak, Aleksandra, Grzegorz Kudla, Tatiana Dudnakova, and David Tollervey. 2013. "Mapping the Human miRNA Interactome by CLASH Reveals Frequent Noncanonical Binding." *Cell* 153 (3): 654–65. https://doi.org/10.1016/j.cell.2013.03.043.

Moore, Michael J., Troels K. H. Scheel, Joseph M. Luna, Christopher Y. Park, John J. Fak, Eiko Nishiuchi, Charles M. Rice, and Robert B. Darnell. 2015. "miRNA-Target Chimeras Reveal miRNA 3'-End Pairing as a Major Determinant of Argonaute Target Specificity." *Nature Communications* 6 (November): 8864. https://doi.org/10.1038/ncomms9864.
